# Mechanisms by which barrier-to-autointegration factor regulates dynamics of nucleocytoplasmic leakage and membrane repair following nuclear envelope rupture

**DOI:** 10.1101/2023.12.21.572811

**Authors:** Charles T. Halfmann, Kelsey L. Scott, Rhiannon M. Sears, Kyle J. Roux

## Abstract

The nuclear envelope (NE) creates a barrier between the cytosol and nucleus during interphase that is key for cellular compartmentalization and protecting genomic DNA. NE rupture can expose genomic DNA to the cytosol and allow admixture of the nuclear and cytosolic constituents, a proposed mechanism of cancer and NE-associated diseases. Barrier-to-autointegration factor (BAF) is a DNA-binding protein that localizes to NE ruptures where it recruits LEM-domain proteins, A-type lamins, and participates in rupture repair. To further reveal the mechanisms by which BAF responds to and aids in repairing NE ruptures, we investigated known properties of BAF including LEM domain binding, lamin binding, compartmentalization, phosphoregulation of DNA binding, and BAF dimerization. We demonstrate that it is the cytosolic population of BAF that functionally repairs NE ruptures, and phosphoregulation of BAF’s DNA-binding that enables its ability to facilitate that repair. Interestingly, BAF’s LEM or lamin binding activity appears dispensable for its role in functional repair. Furthermore, we demonstrate that BAF functions to reduce the extent of leakage though NE ruptures, suggesting that BAF effectively forms a diffusion barrier prior to NE repair. Collectively, these results enhances our knowledge of the mechanisms by which BAF responds to NE ruptures and facilitates their repair.

## Introduction

Interphase compartmentalization of the nucleus in eukaryotic cells is established by the NE, a double membraned extension of the endoplasmic reticulum (ER). Regulated passage of molecules between the nucleus and the cytoplasm occurs via nuclear pore complexes (NPCs) that create holes in the NE. Sequestration of the nucleus from the cytoplasm has several functions, including protection of the genome from cytosolic nucleases, spatial separation of transcription and translation, and enabling a myriad of cell signaling events. In metazoa, the NE is actively disassembled during mitosis and then reformed during mitotic exit, both of which are highly regulated processes. During interphase, the integrity of the NE is protected from mechanically induced trauma by several structures, perhaps most critically the nuclear lamina, a protein meshwork located between the inner nuclear membrane (INM) and the nuclear interior that is composed of type-V intermediate filaments (Burke and Stewart, 2013). Other cellular structures can also contribute to protection of NE integrity. Within the nucleus, the chromatin itself can generate force resisting structures as it integrates with INM proteins and the lamina (Stephens *et al*., 2017). The LINC complex that serves to physically tether the nucleoskeleton to the cytoskeleton sits at the crux of not just force transmission between the cytoplasm and the nucleus, but also may facilitate and/or generate force resistance (Chang *et al*., 2015). Cytoplasmic intermediate filaments often surround the nucleus like a cage and can act as a force absorbing structure within the cytoplasm (Patteson *et al*., 2019). Cell-cell junctions serve to transmit and buffer forces acting on adjacent cells, as does the extracellular matrix.

Despite these various protective mechanisms, the NE can be compromised in a process commonly referred to as nuclear rupture. Confinement and mechanical strain on the nucleus, which can occur during cell migration through a narrow passage, is an example of naturally occurring conditions that can lead to nuclear rupture (Denais *et al*., 2016; Hatch and Hetzer, 2016; Raab *et al*., 2016; Irianto *et al*., 2017). Experimentally, confinement can be induced in vitro, and nuclear rupture can be induced with high curvature probes (Xia *et al*., 2018; Zhang *et al*., 2018; Pfeifer *et al*., 2022) or even laser-induced denaturation of discrete foci on the NE (Denais *et al*., 2016; Halfmann *et al*., 2019; Young *et al*., 2020; Kono *et al*., 2022; Sears and Roux, 2022). Alternatively or in parallel to these approaches to induce nuclear rupture, the structures that protect NE integrity, for example the lamina, can be perturbed making the nucleus susceptible to cellular forces that normally act on the nucleus or experimentally induced forces (Chen *et al*., 2018; Earle *et al*., 2018; Chen *et al*., 2019). Naturally occurring disease-associated mutations in many of these various protective structures, most notably A-type lamins, can lead to weakening of the NE resulting in spontaneous nuclear ruptures (De Vos *et al*., 2011; Tamiello *et al*., 2013). Loss of A-type lamins results in nuclear rupture in myocytes (Earle *et al*., 2018) and progeria-associated mutations in A-type lamins leads to nuclear rupture in vascular smooth muscle cells (Kim *et al*., 2021). Perturbation of nuclear morphology and NE integrity, combined with nuclear confinement during metastasis can lead to nuclear rupture in cancer cells that may contribute to disease pathogenesis (Vargas *et al*., 2012; Denais *et al*., 2016; Yang *et al*., 2017; de Freitas Nader *et al*., 2021).

The reported consequences of nuclear rupture include DNA damage (Denais *et al*., 2016; Raab *et al*., 2016; Earle *et al*., 2018; Xia *et al*., 2018; Chen *et al*., 2019; Patteson *et al*., 2019; de Freitas Nader *et al*., 2021), altered cell proliferation (Cho *et al*., 2019; de Freitas Nader *et al*., 2021) and differentiation (Smith *et al*., 2019). But typically, nuclear rupture does not lead to cell death due to a coordinated repair process that in many ways recapitulates NE reformation following mitosis. Exposure of genomic DNA to the cytosol is the first known trigger that a nuclear rupture has occurred, and it leads to binding of at least two soluble cytoplasmic dsDNA binding proteins, cGAS and barrier to autointegration factor (BAF). As part of the cGAS-STING innate immune response to cytosolic DNA, cGAS stably binds to the dsDNA exposed following rupture and does not rapidly disperse further into the nucleus. Cytosolic BAF competes with cGAS in binding newly exposed genomic DNA following nuclear rupture (Guey et al., 2020), and rapidly moves throughout the nucleoplasm and along the NE. It is capable of binding not only dsDNA, but also proteins bearing the Lap2/Emerin/Man1 (LEM) domain and A-type lamins . BAF is localized to the nucleoplasm, NE and cytoplasm where it serves a variety of functions including protection against dsDNA-based viruses (Suzuki and Craigie, 2002), DNA repair (Bolderson *et al*., 2019), reformation of the postmitotic NE by recruitment of LEM domain proteins and A-type lamins (Haraguchi *et al*., 2001; Margalit *et al*., 2005), and cohesion of chromosomes by DNA crosslinking following mitosis to create a single nucleus (Samwer *et al*., 2017). Post-translational modification of BAF by vaccinia-related kinase 1 (VRK1) and VRK2A phosphorylates BAF on N-terminus threonine and serine residues which inhibits DNA-binding (Nichols *et al*., 2006; Gorjanacz *et al*., 2007a; Molitor and Traktman, 2014; Marcelot *et al*., 2021), and is dephosphorylated by protein phosphatases PP2A, PP4, and the cofactor LEM-domain protein ANKLE2 (Gorjanacz *et al*., 2007a; Asencio *et al*., 2012; Zhuang *et al*., 2014; Mehsen *et al*., 2018). Regulating BAF DNA-binding has also been reported to play roles in suppressing viral DNA in the cytosol (Wiebe and Traktman, 2007; Ibrahim *et al*., 2011; Ibrahim *et al*., 2013; Jamin *et al*., 2014a). We and others have previously shown that BAF is required for recruitment of A-type lamins, LEM domain proteins and their associated-membranes, and ESCRT-III membrane repair machinery to sites of nuclear rupture and is critical to the functional repair of nuclear ruptures (Halfmann *et al*., 2019; Young *et al*., 2020; Kono *et al*., 2022). Here, we follow up on our previous studies exploring the functional role of BAF in the response to nuclear rupture and the process of NE repair.

## Results

### Development of a new NE rupture reporter that is nucleo-cytoplasmic transport-independent

In our previous study, we utilized GFP-BAF to observe how mutations that perturb specific functions with DNA and LEM-domain proteins impacted dynamics before and during NE rupture. Here, we sought to further examine how these BAF variants impacted the functional role of NE rupture repair. Unfortunately, our initial experiments with NIH3T3 mouse fibroblasts indicated that GFP-BAF was unable to functionally rescue NE rupture repair upon depletion of endogenous BAF with siRNAs (Supplemental Figure S1). In light of these findings, we stably expressed variants of an siRNA-resistant BAF while simultaneously co-expressing a nuclear rupture reporter using an IRES bicistronic vector in BJ5ta human diploid fibroblasts (Figure 1A-C and 1F-H). These cells were used to assess the ability of the BAF variants to control the extent of rupture leakage and functionally repair nuclear ruptures. Most previous studies of nuclear rupture have utilized a rupture reporter composed of a GFP fused to a nuclear localization sequence (GFP-NLS) that during interphase is predominantly nuclear due to constant reimport of the GFP as it passively diffuses out of the nucleus through the NPCs. Upon nuclear rupture, the GFP-NLS leaks out into the cytoplasm at a rate that import is unable to overcome, but regains nuclear localization upon sufficient NE repair as the rate of nuclear import exceeds the extent of leakage into the cytoplasm (Figure 1D and 1E). NE repair can be verified by photobleaching residual GFP-NLS in the cytoplasm after rupture and monitoring for any continued nuclear GFP-NLS diffusing into the cytoplasm (Figure 1D). Given recent evidence that changes in the NE constituency and/or changes in cellular forces can significantly alter nuclear transport (Andreu et al., 2022; Scott *et al*. manuscript in preparation), we developed a rupture reporter that functions independent of facilitated nucleocytoplasmic transport. Fusion of GFP to the large tetrameric Hsp90 (Hsp90-GFP; Figure 1F-H) leads to predominant exclusion of the GFP from the interphase nucleus due to its size, which considerably exceeds the diffusion barrier limit of NPCs. Upon NE rupture, Hsp90-GFP diffuses through the site of rupture into the nucleus until the barrier is resealed (Figure 1J). Subsequent photobleaching of the nuclear Hsp90-GFP can be used to assess the extent of restoration of NE barrier function, which we term a repair check (Figure 1I). Throughout most of the studies here, GFP-NLS was used in parallel to HSP90-GFP to validate this new rupture reporter. For both reporters, the initial rate of diffusion into the restricted compartment can be used as a readout to examine the extent of rupture, which can be affected by the size of the rupture and any diffusion barrier that may exist (Figure 1E and 1J). For these studies we utilized laser-induced nuclear rupturing, a method that applies a brief pulse of high intensity 405 nm light exposure at a specific site on the periphery of the NE to denature the proteins that create the structural scaffold that maintains NE membrane integrity, which likely includes the lamina, LINC complex, NPCs and peripheral heterochromatin. At variable times following laser ablation but typically within 30 sec (Supplemental Figure S2), the NE ruptures, presumably due to focal weakening of structures that support the NE such that they are unable to counterbalance intrinsic forces within the cell that act on the nuclear membranes. To assess the relationship between NPCs and sites of laser-induced NE rupture, we used the highly dynamic 3GFP-Nup153 as an NPC marker to overcome photobleaching of the more stable NPC constituents. Interestingly, we observed initial accumulation of rupture reporters at NPCs that quickly spread into discrete, NPC-free regions on the NE (Supplemental Figure S3). We also observed a loss of 3GFP-Nup153 from the rupture sites, likely due to a denaturation of the NPC structure. This could possibly indicate that the loss of structural integrity on the NE could initiate from NPCs. Although artificial, this laser-induced NE rupture method generates a temporospatially consistent rupture event that allows for more reliable quantification of loss and repair of NE barrier function, including the rate of leakage of rupture reporters and more precise timing of repair. NE rupture can be verified by the nuclear rupture marker cGAS-mCherry localizing to the site of rupture, as well as the rapid loss of nuclear GFP-NLS or nuclear accumulation of HSP90-GFP.

**Figure 1:**
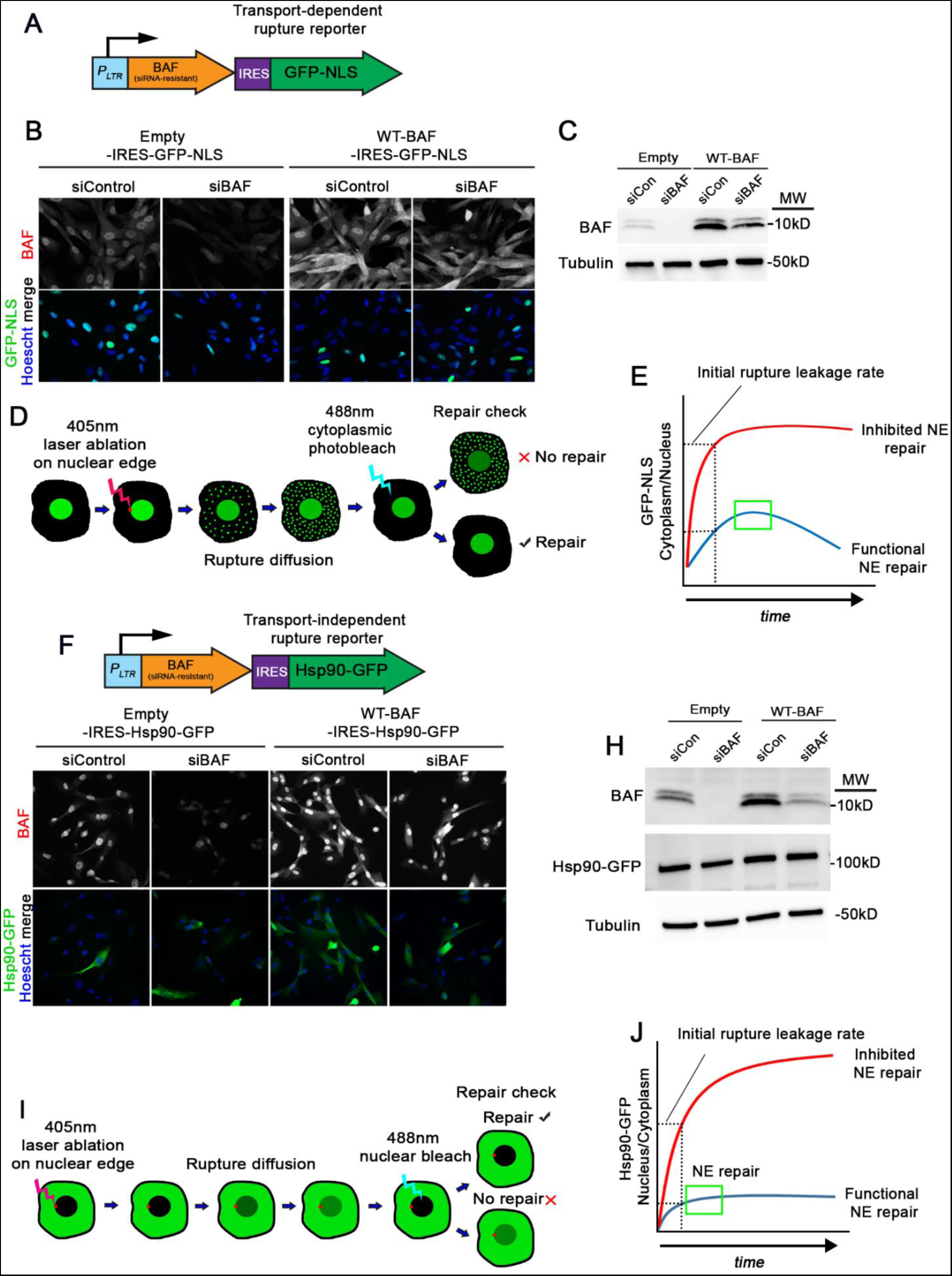
Development of a nuclear transport-independent NE rupture reporter. A) Illustration of the transport-dependent BAF-IRES-GFP-NLS rupture reporter system for co-expression with BAF variants. P_LTR_, long terminal repeat promoter. B) IF images of BJ5ta cells expressing either empty vector- or WT-BAF-IRES-GFP-NLS probed with BAF antibody after treatment with siControl or siBAF. C) Representative western blots of cell lysates from cells in A probed with a BAF antibody. MW; molecular weight. D) Illustration of workflow for testing GFP-NLS diffusion and repair of NE ruptures in cells expressing BAF variants. E) Graphical representation of measuring early rupture diffusion and repair (blue line) or inhibited repair (red line) in cells from D. F) Illustration of the transport-independent BAF-IRES-Hsp90-GFP rupture reporter system for co-expression of BAF variants, along with IF images (F), and western blots (H). I) Illustration of workflow for testing Hsp90-GFP rupture diffusion and repair, as well as graphical representation of Hsp90-GFP rupture curves in cells expressing BAF variants. Scale bar; 10 µm.

### Endogenous levels of BAF enable optimal efficiency in regulating the extent of NE rupture and repair

We first used the GFP-NLS and Hsp90-GFP rupture reporters to explore how overexpression of a siRNA-resistant wild-type BAF (WT-BAF) controlled the extent and repair of NE ruptures. In cells expressing empty vector-IRES-Hsp90-GFP (Empty) transfected with control siRNAs (siControl), a nominal amount of Hsp90-GFP diffused into the nucleus following rupture (Figure 2A) and leveled off after roughly 1-2 minutes (Figure 2B). Depletion of endogenous BAF resulted in a much larger extent of Hsp90-GFP diffusion into the nucleus (Figure 2A and 2B) which saturated the nucleus within 5 minutes (Figure 2A). Cells overexpressing the siRNA-resistant WT-BAF, treated with either siControl or siBAF, had similar Hsp90-GFP diffusion into the nucleus post-rupture as untreated cells with endogenous BAF (Figure 2B). Additionally, the rate of initial Hsp90-GFP diffusion was similar between cells with endogenous BAF compared to cells with elevated WT-BAF, in contrast with cells lacking BAF (Figure 2C). Repair checks in cells expressing endogenous or WT-BAF revealed functional repair of the NE barrier within 10 min after rupture, whereas many BAF-depleted cells lacked a functional NE barrier (Figure 2D). These results reveal that endogenous levels of BAF are sufficient for repair of NE ruptures and elevating the level of BAF does not appreciably slow rupture leakage or shorten the duration of NE repair. We observed similar results utilizing cells expressing GFP-NLS instead of HSP90-GFP (Supplemental Figure S4A), supporting that the new transport independent reporter is functioning as intended.

**Figure 2:**
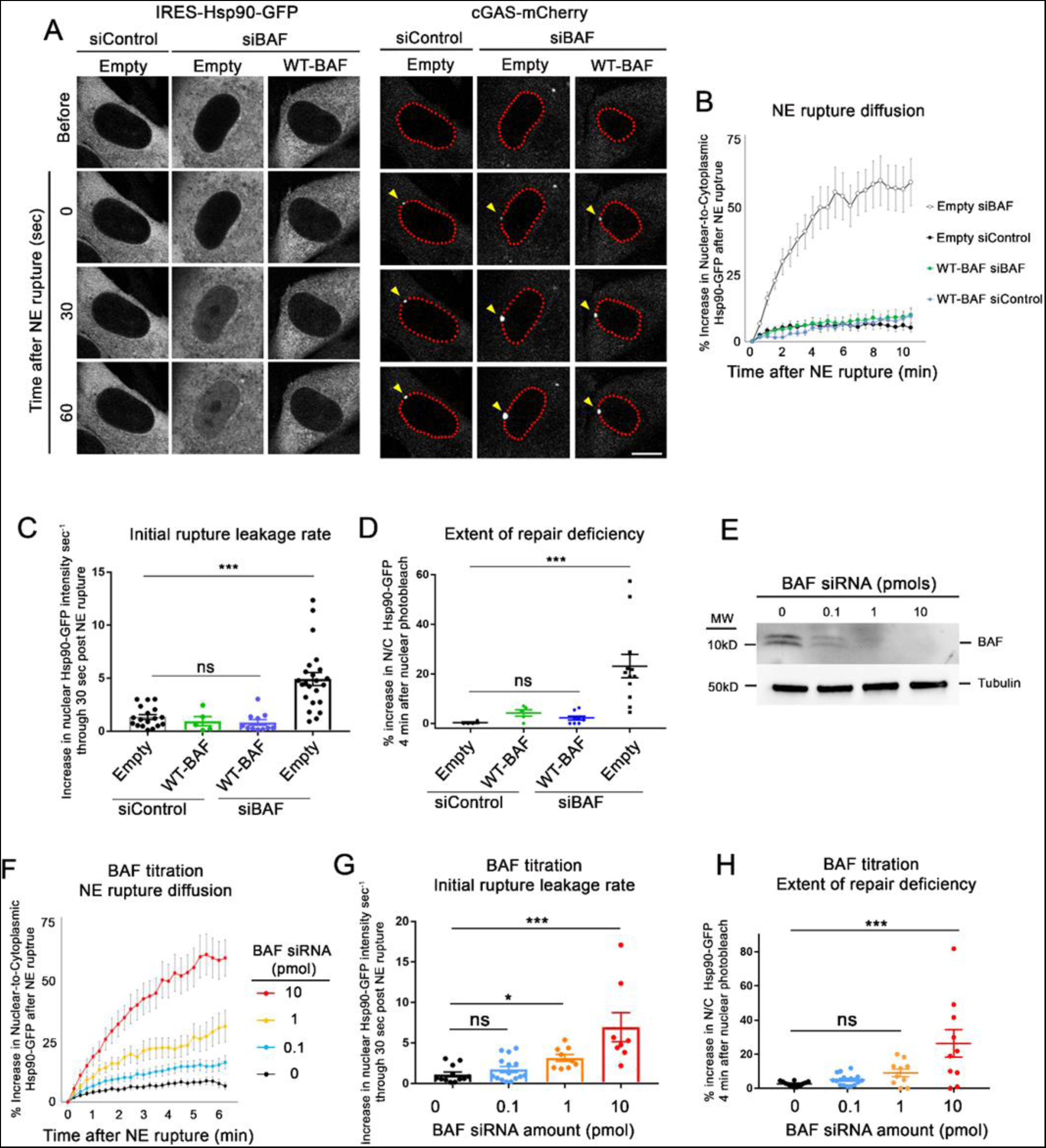
Endogenous BAF is sufficient to respond to NE ruptures with gradual depletion leading to a progressive loss of that response. A) Representative images of BJ5ta cells expressing either Empty- or WT-BAF-IRES-Hsp90-GFP and cGAS-mCherry, treated with either siControl or siBAF for 96 hr and followed through NE rupture to observe the extent of leakage. The enrichment of cGAS (yellow arrows) along the nuclear periphery (outlined in red for clarity) signal the location and time of NE rupture. Scale bar, 10 µm. B) Quantification of the nuclear-to-cytoplasmic increase in Hsp90-GFP following laser-induced NE rupture from cells. The graph represents mean values ± SEM of Empty siControl (n= 18), Empty siBAF (n=18), WT-BAF siControl (n=5) and WT-BAF siBAF (n=11) cells from at least three independent experiments. C) Rates of Hsp90-GFP leakage into the nucleus during the first 30 seconds following NE rupture. D) Repair check, measured as a percent increase in nuclear-to-cytoplasmic Hsp90-GFP diffusion into the nucleus following nuclear photobleaching (performed 10 minutes after NE rupture) to check functional repair. The graph represents mean values ± SEM of Empty siControl (n = 4), WT-BAF siControl (n = 5), WT-BAF siBAF (n = 8) and Empty siBAF (n = 10). E) Representative Western blot of cell lysates from cells treated with 0, 0.1, 1 and 10 pmol of BAF siRNAs for 72 hr, with tubulin used as a loading control. F) Quantification of the nuclear-to cytoplasmic increase in Hsp90-GFP following laser-induced NE rupture from cells treated with 0, 0.1, 1 and 10 pmol of BAF siRNAs (n= 12, 14, 9, and 11 cells, respectively) from at least two independent experiments. G) Initial rates of Hsp90-GFP leakage into the nucleus during the first 30 seconds following NE rupture from E. H) Repair check of cells from F, 10 min post-rupture. Statistical significance: *, P<0.05; **<0.005; ***, P<0.0005 using a one way ANOVA with Dunnett’s post hoc multiple comparison test.

Since we observed no change in the extent of NE rupture or rate of repair with elevated levels of BAF, we next investigated if a progressive reduction of BAF led to a progressive change in these processes or if there was an abrupt change upon loss of a critical amount of BAF. We measured the extent of Hsp90-GFP leakage following NE rupture when BAF levels were incrementally reduced by titration of BAF siRNA. Western blots confirmed a gradient of cellular BAF loss in cells treated with increasingly diluted siBAF (Figure 2E), and the extent of Hsp90-GFP leakage following NE rupture increased as BAF levels were incrementally lowered (Figure 2G and 2H), including a gradual increase in the number of cells that failed to repair their NE rupture (Fig 2H). These results suggest that a reduction of BAF correlates with a gradual increase of rupture leakage and decrease in repair efficiency.

### BAF must be cytosolic to facilitate repair of NE rupture

Our previous studies using compartmentalized photobleaching of GFP-BAF, which does not appreciably exchange between the cytoplasm and nucleus, suggested that it is predominantly the cytoplasmic population of BAF that localizes to sites of NE rupture (Halfmann *et al*., 2019). Here we investigated whether the functional repair of NE ruptures is dependent on the cytoplasmic and/or nuclear pool of BAF. To exclude BAF from either the nucleus or cytosol, we fused an N-terminal nuclear export sequence (NES) or an NLS to WT-BAF (NES-BAF and NLS-BAF, respectively) and confirmed appropriate localization by immunofluorescence microscopy (Figure 3A, Supplemental Figure S4B). With GFP-NLS as a reporter, we observed similar rupture leakage in cells depleted of endogenous BAF and expressing NES-BAF or WT-BAF (Supplemental Figure S4C). With Hsp90-GFP as a reporter we observed that diffusion into the nucleus was slightly elevated in cells depleted of endogenous BAF and expressing NES-BAF, as compared to those expressing WT-BAF, although expression of NES-BAF led to a similar initial rupture leakage rate of Hsp90 upon rupture and was capable of repairing NE ruptures within 10 minutes (Figure 3C). In contrast, NLS-BAF was incapable of repairing NE ruptures with either the GFP-NLS (Supplemental Figure S4C) or Hsp90-GFP reporter (Figure 3C), and this was accompanied by a considerable increase in the initial rate of reporter leakage (Figure 3C). These results support the hypothesis that it is the cytoplasmic BAF that prevents substantial leakage following NE rupture and facilitates repair of NE ruptures, whereas nuclear BAF does not appear to participate in these processes.

**Figure 3:**
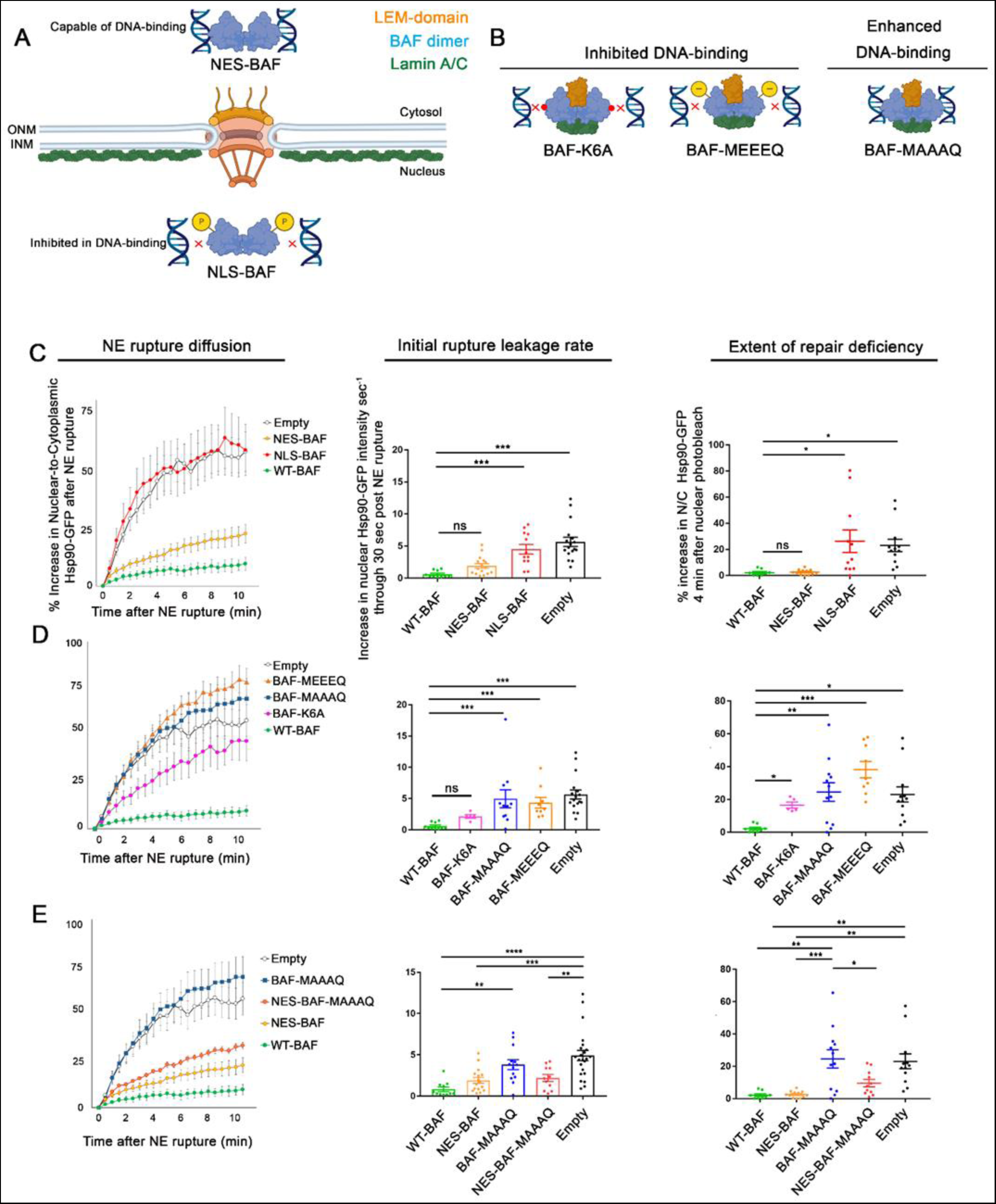
Cytosolic localization and phosphoregulation of DNA binding is critical to BAF’s response to NE rupture. A) Design of variations predicted to alter BAF’s cellular localization in the cytosol and nucleus (NES- and NLS-BAF, respectively), which influences DNA-binding via phosphorylation. B) DNA-binding BAF variants predicted to inhibit (BAF-K6A and BAF-MEEEQ) or enhance (BAF-MAAAQ) DNA-binding. Illustrations show BAF dimer (blue) in relationship to LEM-domain (orange) and lamin A/C (green) binding sites. C) Quantification of the nuclear-to-cytoplasmic ratio of Hsp90-GFP after NE rupture in cells expressing Empty-, WT-BAF, NES-BAF, and NLS-BAF (n = 17, 10, 16, and 12 cells respectively), with initial rates of Hsp90-GFP diffusion into the nucleus and repair efficiency of NE ruptures, measured by the increase of nuclear Hsp90-GFP after nuclear photobleaching 10 min following rupture (n = 17, 10, 16, and 12 cells respectively). D) Quantification of the nuclear-to-cytoplasmic ratio of Hsp90-GFP after NE rupture in cells expressing BAF variants with inhibited DNA binding (BAF-MEEEQ and BAF-K6A; n = 9 and n = 5, respectively) or enhanced DNA-binding (BAF-MAAAQ; n = 14), with initial rate of Hsp90-GFP increase into the nucleus after NE rupture and repair efficiency measurements of NE ruptures through nuclear photobleaching 10 min after NE rupture. E) Quantification of the nuclear-to-cytoplasmic ratio of Hsp90-GFP after NE rupture in cells expressing Empty-, WT-BAF BAF-MAAAQ, NES-BAF, and NES-MAAAQ BAF (NES-MAAAQ; n = 11), with initial rate of Hsp90-GFP increase into the nucleus following NE rupture of cells, and repair efficiency of NE ruptures using nuclear photobleaching. Statistical significance: *, P<0.05; **<0.005; ***, P<0.0005; Figures C-D using a one way ANOVA with Dunnett’s post hoc multiple comparison test, Figures E using a one way ANOVA with Tukey’s post hoc multiple comparison test.

### The phosphoregulation of BAF’s DNA binding is critical to its response to NE rupture

The differences in BAF’s ability to respond to NE ruptures based on its cytoplasmic or nuclear localization is likely due to its differential phosphorylation within those two compartments. Nuclear BAF, in close proximity to its kinases VRK1 and VRK2, is highly phosphorylated at its N-terminus that reduces affinity to DNA (Nichols *et al*., 2006; Marcelot *et al*., 2021), and therefore would be less able to bind DNA at a rupture site. Furthermore, there would be no trigger for nuclear BAF to bind to DNA exposed at the rupture site, as any nuclear BAF that was capable of binding DNA would already be pre-bound to DNA throughout the nucleus. In contrast, the retention of BAF in the cytoplasm, away from its nuclear kinases and closer proximity to the phosphatases that regulate it, such as PP4 and PP2A (Asencio *et al*., 2012; Gorjánácz, 2013), is likely unphosphorylated and primed to bind DNA, much as it is known to bind cytoplasmic viral DNA (Chen and Engelman, 1998; Lee and Craigie, 1998; Wiebe and Traktman, 2007; Ibrahim *et al*., 2013; Jamin *et al*., 2014b) or even exogenous DNA (Kobayashi *et al*., 2015; Burger *et al*., 2020). To test how BAF phosphoregulation affects NE rupture leakage and repair, BAF’s N-terminal phosphorylates residues (M<u>TTS</u>Q) were replaced with either alanine (MAAAQ) or glutamic acid (MEEEQ) to prevent or mimic phosphorylation, respectively (Figure 3B). As has been reported (Jamin *et al*., 2014a; Jamin *et al*., 2014b; Halfmann *et al*., 2019), prevention of BAF phosphorylation resulted in a substantial shift of BAF from the cytosol to the nucleus (Supplemental Figure S4E), likely due to enhanced DNA binding. Expectedly, based on our results with the NLS-BAF, cells with BAF-MAAAQ exhibited an enhanced initial rate of reporter leakage (Figure 3D and Supplemental Figure S4F) and an inability to efficiently repair NE ruptures (Figure 3D). We hypothesized that the repair defect most likely reflects the loss of BAF from the cytoplasm due to being in a constant state of high affinity DNA-binding, thus it is unable to target to and enrich at NE ruptures. To test this, we fused an NES to BAF-MAAAQ (NES-BAF-MAAAQ) to drive it into the cytoplasm, where we hypothesized it would be able to rescue the loss of endogenous BAF. Indeed, we observed that expression of NES-BAF-MAAAQ reduced Hsp90-GFP rupture leakage and improved NE repair compared to BAF-MAAAQ (Figure 3E), suggesting that the repair defect observed in BAF-MAAAQ is due to an inability to access DNA exposed to the cytosol at rupture. In contrast, the phosphomimetic BAF-MEEEQ is localized in both the cytosolic and nuclear compartments (Supplemental Figure S4E) but led to a similarly large initial rate of reporter leakage and an inability to repair the NE barrier as a total loss of BAF (Figure 3D). This is likely due to the profoundly reduced affinity to DNA, since a BAF variant inhibited in DNA binding (BAF-K6A) also exhibited similar defects with both rupture reporters (Figure 3D, Supplemental Figure S4D). These results suggest that the compartmental phosphoregulation of BAF is critical to facilitate repair of NE ruptures.

### BAF homodimerization is requisite for efficient repair of NE rupture

Due to its ability to homodimerize and perhaps further oligomerize, BAF has the ability to condense DNA (Zheng *et al*., 2000; Skoko *et al*., 2009). Previously, this function was reported to be critical to NE reformation where, upon mitotic exit, BAF binds to and crosslinks DNA into a stiff chromatin network that aids in membrane reformation around the nucleus (Samwer *et al*., 2017). In order to understand how BAF-mediated DNA condensation effects the nuclear leakage and repair of ruptures, we utilized a previously known G47E mutation that impairs BAF homodimerization, but does not prevent DNA-binding (Segura-Totten *et al*., 2002). Since BAF-G47E has previously been shown to be retained in the nucleus (Ibrahim *et al*., 2011), likely due to loss of phosphorylation and thus unregulated DNA binding, we fused an NES to the N-terminus (NES-BAF-G47E) to increase its presence in the cytoplasm and ability to bind to NE ruptures (Figure 4A, Supplemental Figure 5A). To assess NES-BAF-G47E localization at sites of NE rupture, we fixed cells shortly after induced rupture and immunolabeled BAF. Within 10 seconds after NE rupture, both WT- and NES-BAF-G47E are similarly enriched at rupture sites, although by 4 minutes post-rupture levels of NES-BAF-G47E were more diminished compared to WT-BAF or NES-BAF (Figure 4B), suggesting that the loss of BAF dimerization prevents retention at the rupture site. NES-BAF-G47E also inhibited emerin recruitment to ruptures as expected (Figure 4C), since the LEM-domain binding site sits at an interface between BAF dimers (Figure 4A). Compared to NES-BAF or WT-BAF, NES-BAF-G47E was unable to fully rescue the initial rate of reporter leakage or functional NE rupture repair (Figure 4D-F, Supplemental Figure 5B), potentially due in part to an inability to condense DNA and/or to interact with LEM domain proteins and/or A-type lamins that bind to the dimer.

**Figure 4:**
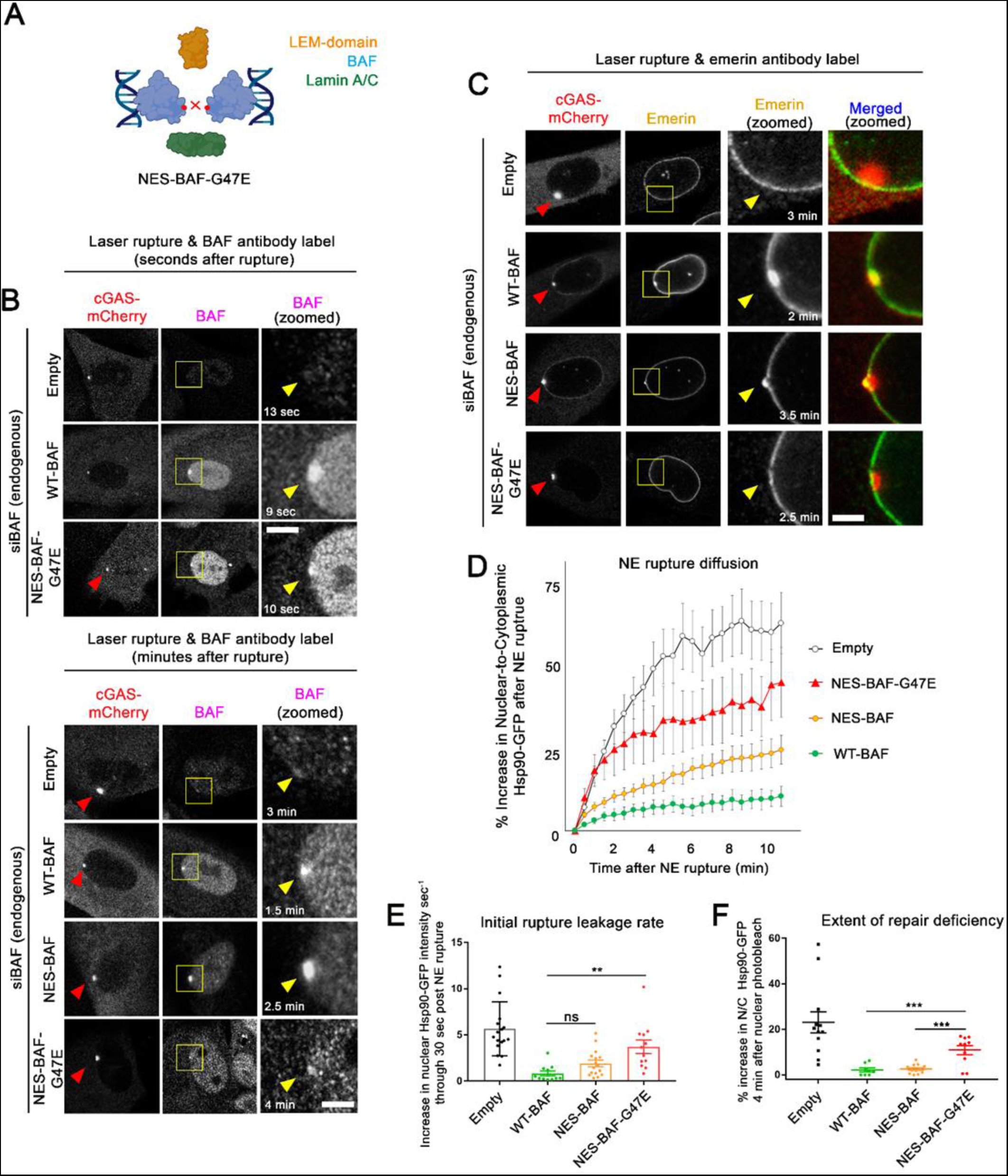
BAF homodimerization is critical for controlling rupture leakage and the repair of NE ruptures. A) Model of NES-BAF-G47E monomers effect on homodimerization, and protein and DNA interactions. B) Representative images of cells expressing BAF variants treated with siBAF, then fixed and labeled with BAF antibody at indicated time points following laser-induced NE rupture. Scale bar, 2µm. Images are representative of the following number of cells: Empty, n=4; WT-BAF, n=4; NES-BAF, n=5; NES-BAF-G47E, n=5. C) Representative images of cells expressing BAF variants treated with siBAF, then fixed and labeled with Emerin antibody at indicated time points following laser-induced NE rupture. Images are representative of the following number of cells: Empty, n=4, WT-BAF, n=2; NES-BAF, n=5, NES-BAF-G47E, n=4 Scale, 2µm. D) Quantification of the nuclear-to-cytoplasmic ratio of Hsp90-GFP after NE rupture in cells expressing Empty-, WT-BAF, NES-BAF, and NES-BAF-G47E (n=17, 10, 16, and 12 cells respectively) from at least duplicate experiments with initial rate of Hsp90-GFP increase into the nucleus following NE rupture of cells (E) and repair efficiency of NE ruptures using nuclear photobleaching (F). Error bars indicate ± SEM. Statistical significance: *, P<0.05; **<0.005; ***, P<0.0005 using a one way ANOVA with Tukey’s post hoc multiple comparison between WT-BAF, NES-BAF, and NES-BAF-G47E. Empty shown for relative comparison.

### LEM-domain or lamin interaction with BAF are dispensable for functional repair of NE rupture

LEM-domain proteins have been shown to be dependent on BAF for their recruitment to NE ruptures (Halfmann *et al*., 2019; Young *et al*., 2020; Janssen *et al*., 2022a). A-type lamins, which interact with BAF via association with lamin’s IgG-like β-fold domain, are also rapidly recruited and stabilized to NE ruptures in a BAF-dependent manner (Young *et al*., 2020, Kono *et al*., 2022; Sears and Roux, 2022). Loss of LEM-domain proteins to NE ruptures are reported to have deleterious effects on the repair of NE ruptures. A combinatorial loss of multiple transmembrane LEM-domain proteins (Emerin, LEMD2, and Ankle2) results in a failure to facilitate efficient NE repair (Halfmann *et al*., 2019), and expression of a mutated GFP-BAF lacking a LEM-domain binding site (L58R) was reported to target to NE ruptures but increased the duration of NE repair (Young *et al*., 2020). A progeria-associated BAF-A12T mutation that disrupts BAF-lamin binding in Nestor-Guillermo Progeria Syndrome (NGPS) been reported to inhibit emerin and lamin recruitment to nuclear blebs and increased occurrence of re-rupturing events (Janssen *et al*., 2022a). These findings suggest that LEM-domain proteins and lamins play roles in functional NE rupture repair by recruiting membrane and, in the case of LEMD2, ESCRT-III membrane repair machinery; or in the case of lamins, stabilizing the rupture site to protect against re-rupturing. To examine the functional role of BAF-recruited LEM-domain proteins and lamins at sites of NE rupture, we assessed how the well-characterized L58R and A12T BAF variants impact the rate of NE rupture leakage and repair (Figure 5A, Supplemental Figure 5C). NE ruptured cells depleted of endogenous BAF and expressing the siRNA-resistant BAF variants were fixed within 4 minutes of cGAS-mCherry enrichment at rupture sites and probed with specific antibodies to verify the presence or absence of the LEM-domain protein emerin or A-type lamins (phosphorylated LaA/C; pLaA/C) at sites of NE rupture. Expectedly, both BAF-L58R and BAF-A12T were able to enrich at NE ruptures similar to WT-BAF within 2.5 min of rupture (Fig 5B), in agreement with previous studies that the loss of lamin or LEM-domain binding does not prevent the recruitment of BAF to ruptures (Young *et al*., 2020; Janssen *et al*., 2022a). Cells expressing WT-BAF and BAF-A12T also had a pronounced accumulation of emerin at NE ruptures, whereas cells expressing BAF-L58R displayed a noticeable loss of emerin from the NE and did not enrich emerin at ruptures (Figure 5C), suggesting that the L58R mutation indeed inhibits their recruitment to rupture sites. Likewise, cells expressing WT-BAF and BAF-L58R enriched pLaA/C at ruptures, while cells expressing BAF-A12T did not (Figure 5D). Functionally, both BAF-A12T and BAF-L58R cells displayed similar initial rates of Hsp90-GFP rupture reporter leakage compared to WT-BAF cells and appeared as capable as WT-BAF in repairing ruptures (Figure 5E), suggesting that lamins or LEM-domain proteins are dispensable for controlling rupture leakage and facilitating repair. Similarly, cells expressing BAF-A12T or BAF-L58R also showed GFP-NLS rupture leakage and repair kinetics similar to a WT-BAF expressing cells, although cells expressing BAF-L58R exhibited a slight delay in re-import of the GFP-NLS post-rupture (Supplemental Figure S5D). These results with BAF-A12T are in agreement with a recent study showing that ruptures from NGPS patient cell lines using a GFP-NLS reporter exhibited similar rupture kinetics as control cells (Janssen *et al*., 2022a). However, our findings that inhibiting LEM-domain recruitment has a nominal effect on controlling leakage and facilitating repair was in contradiction to earlier findings demonstrating that a GFP-BAF-L58R was unable to repair NE ruptures (Young *et al*., 2020), or our previous studies showing that a combinatorial loss of multiple LEM-domain proteins caused a failure to repair NE ruptures (see discussion below). Together, these results suggest that BAF enrichment to ruptures in the absence of lamins and LEM-domain protein recruitment is capable of controlling rupture leakage and facilitating repair.

**Figure 5:**
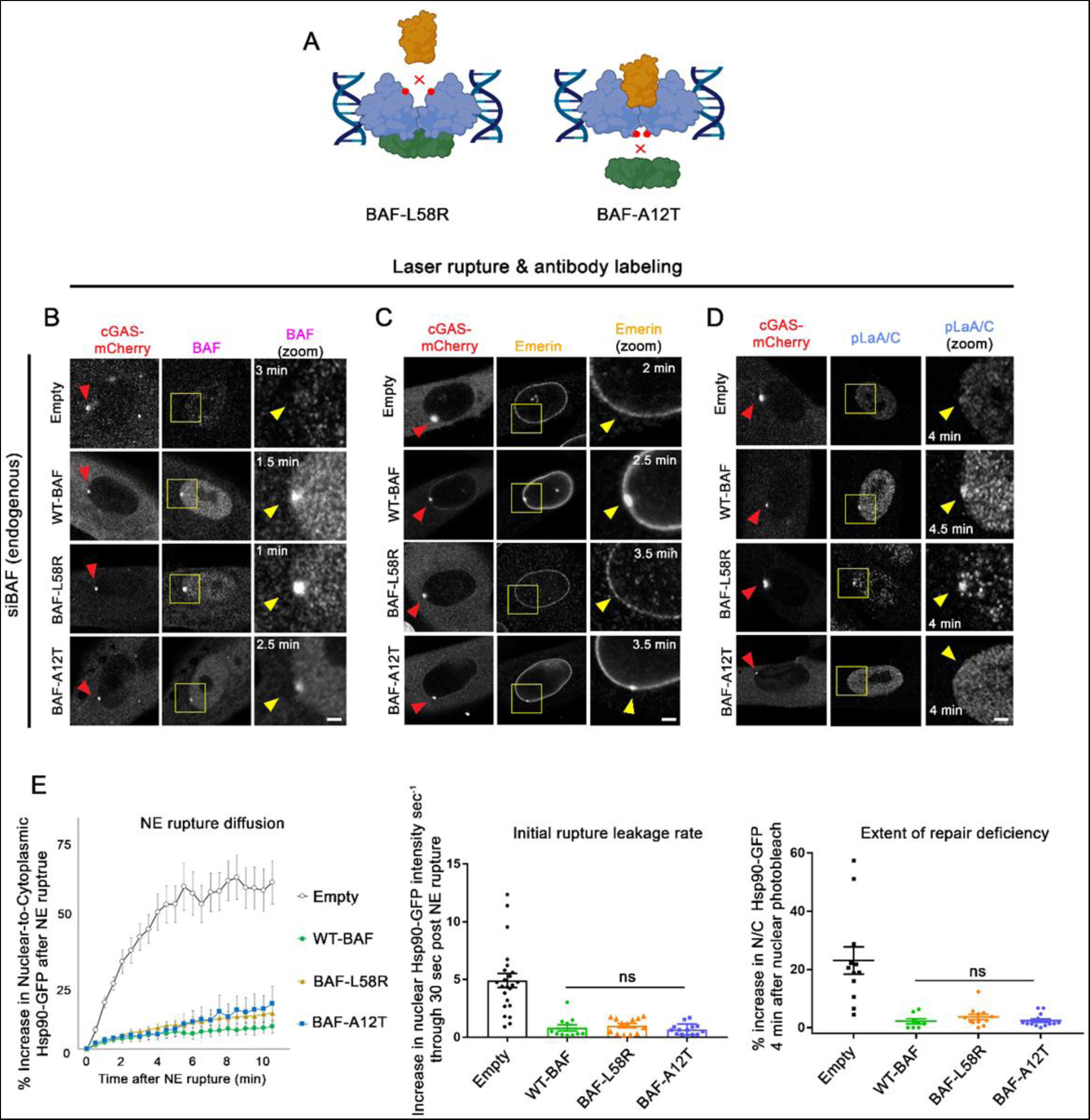
BAF-LEM domain protein and lamin interactions have a negligible impact on the control of leakage and repair of NE ruptures. A) Model of variants predicted to disrupt BAF binding to LEM-domain proteins (L58R) and Lamin A/C (A12T). B) Representative images of cells expressing BAF-WT, -L58R, and -A12T variants treated with siBAF, then fixed and labeled with BAF (B), Emerin (C), and phospho-lamin A/C (D; pLaA/C) antibody at indicated time points following laser-induced NE rupture. Yellow arrow indicates the site of rupture. Scale bar, 2µm. Each image is a representative of the following number of cells. BAF antibody: Empty, n=4; WT-BAF, n=4; BAF-L58R, n=5; BAF-A12T, n=3. Emerin antibody: Empty, n=4, WT-BAF, n=2; BAF-L58R, n=7; BAF-A12T, n=2. pLaA/C antibody: Empty, n=3, WT-BAF, n=5; BAF-L58R, n=2, BAF-A12T, n=5. E) Quantification of the nuclear-to-cytoplasmic ratio of Hsp90-GFP after NE rupture in Empty-, WT-BAF, BAF-L58R, and BAF-A12T variants (n=17, 10, 13, and 15 cells respectively) from triplicate experiments, with initial rate of Hsp90-GFP increase into the nucleus following NE rupture and repair efficiency of NE ruptures using nuclear photobleaching. Error bars indicate ± SEM. Statistical significance: *, P<0.05; **<0.005; ***, P<0.0005 using a one way ANOVA with Tukey’s post hoc multiple comparison between WT-BAF, BAF-L58R, and BAF-A12T. Empty shown for relative comparison.

### BAF controls rupture diffusion in a size-dependent manner

During the course of these studies, we observed a considerable increase in the initial rate of Hsp90-GFP diffusion into the ruptured nucleus when BAF was either absent or was incapable of binding and/or sufficiently cross-bridging DNA. We hypothesized multiple factors that could explain this phenomenon. The loss of BAF could somehow lead to a larger hole in the NE following laser-induced rupture. To assess this possibility, we utilized BJ5ta cells expressing a GFP-tagged transmembrane ER protein (GFP-Sec61β) that enables visualization of the NE and measured the width of the NE gap during the first 30 seconds following rupture (verified by cGAS-mCherry). Within 5 seconds after NE rupture, the width of the NE gap was on average ∼26% larger than control cells in BAF-depleted cells and remained open well past 30 sec post-rupture, unlike the progressive membrane closure observed in control cells (Figure 6A, Supplemental Figure S6). It is also possible that the rapid BAF-mediated membrane closure of a nuclear rupture could itself be contributing to the difference in the initial rate of post-rupture leakage. An alternate mechanism by which BAF could slow the leakage of Hsp90-GFP into the nucleus could be its ability to create a diffusion barrier at the rupture site. BAF can condense DNA and was shown to bind the surface of chromatin and create a diffusion barrier to large macromolecules, a process proposed to be critical during post-mitotic NE reformation (Samwer *et al*., 2017). To investigate if BAF has a similar role in creating a diffusion barrier at NE ruptures, we compared the diffusion into the nucleus of Hsp90-GFP to the smaller GFP-α tubulin. The α-β tubulin complex creates a cylindrical structure (length x width, ∼9 nm x 5 nm) that is considerably smaller than the dimeric (length x width, ∼14 nm x 7 nm) or hexameric (length x width, ∼14nm x 14nm) Hsp90 complex. Within the first 30 seconds of NE rupture, GFP-α-tubulin was observed to diffuse into the nucleus at a considerably faster rate than Hsp90-GFP in the presence of BAF (Figure 6B-D). In the absence of BAF, GFP-α-tubulin and Hsp90-GFP exhibited similar rupture diffusion curves (Figure 6C), although initial rupture leakage was slightly higher for GFP-α-tubulin than Hsp90-GFP (Figure 6D). To asses differential mobility of these two proteins in the cytoplasm, we utilized fluorescence loss after photobleaching (FLIP) and found that GFP-α-tubulin exhibited significantly higher mobility than Hsp90 (Figure 6E-G). After normalizing to this cytoplasmic mobility, we found that GFP-α-tubulin had a significantly higher rate of diffusion into the nucleus than Hsp90-GFP when BAF was present, but that both proteins had similar diffusion rates when BAF was depleted (Figure 6H). This data suggests that upon DNA binding at the rupture, BAF is capable of regulating the diffusion of cytosolic proteins into the nucleus, possibly by forming a size-exclusion barrier at the rupture site, and in the absence of this barrier, proteins diffuse through the rupture at similar rates regardless of their relative size. This barrier could be formed by BAF binding DNA and possibly other proteins to create a diffusion barrier, facilitating rapid membrane closure, or a combination of both these processes.

**Figure 6:**
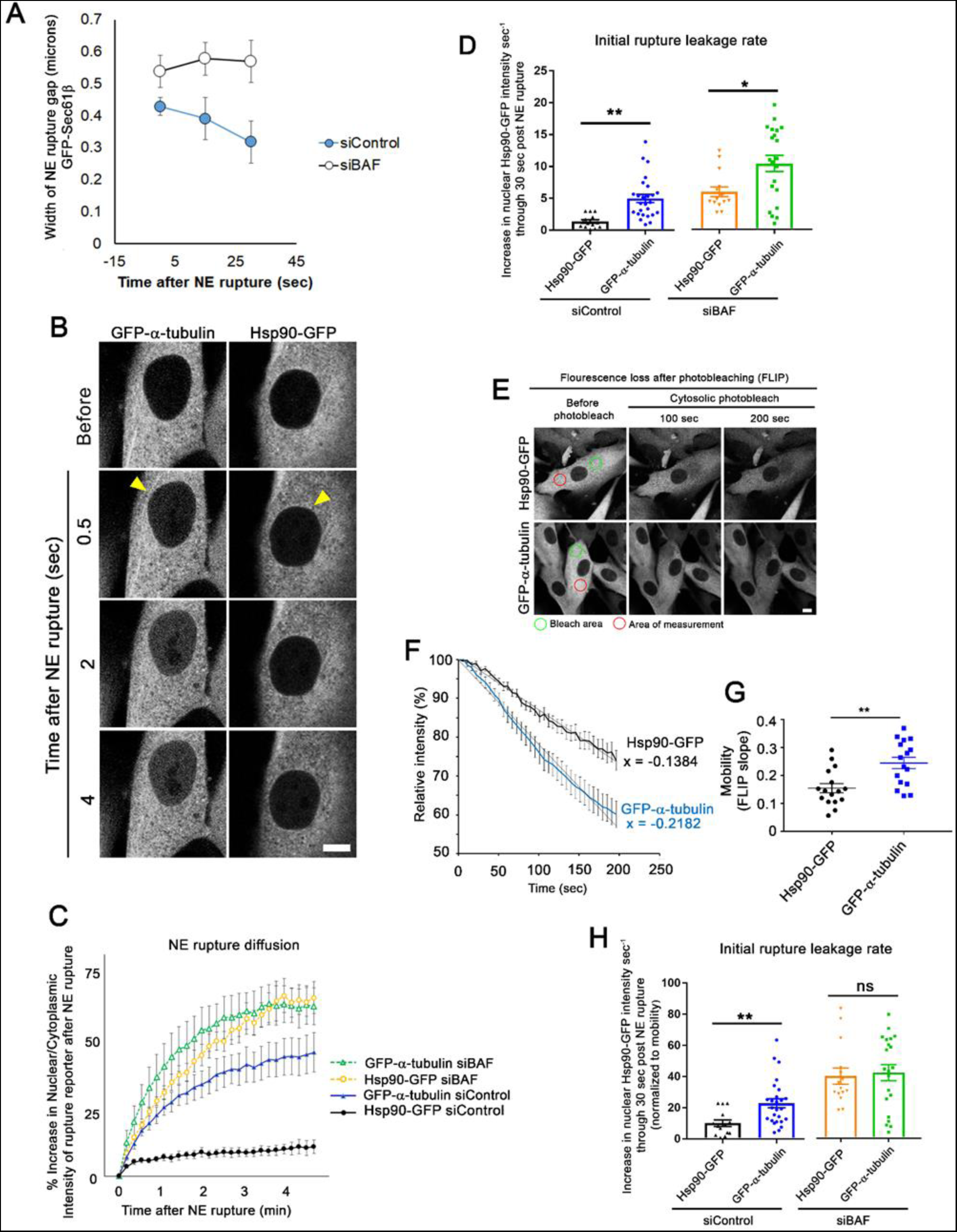
BAF controls rupture diffusion in a size-dependent manner. A) Measurements of the width of the NE rupture gap in BJ5ta cells expressing GFP-Sec61β. Number of cells analyzed: siControl, n=15; siBAF, n=14. Error bars indicate ± SEM from triplicate experiments. B) Representative images of BJ5ta cells expressing either Hsp90-GFP or α-Tubulin-GFP after laser-induced NE rupture. Scale bar; 10 µm. C) Quantification of the nuclear-to-cytoplasmic ratio of cells expressing either Hsp90-GFP or α-tubulin-GFP treated with either siControl (n = 29 and 14, respectively) or siBAF (n = 21 and 16, respectively) from triplicate experiments. Error bars indicate ± SEM. D) Initial rate of increase into the nucleus following NE rupture for cells in B. E) Representative images of BJ5ta cells expressing Hsp90-GFP or α-tubulin-GFP during FLIP. Green circle indicates area of photobleaching in the cytoplasm. Red circle indicates area of measurement. F) FLIP measurements showing relative mobility of Hsp90-GFP (n = 16 cells) or α-tubulin-GFP (n = 16 cells). The slope of each line *x* was used to factor relative mobility between Hsp90-GFP and α-tubulin-GFP. G) Mobility of Hsp90-GFP and α-tubulin-GFP (measured as the best line-of-fit slope for each individual cell). H) Initial rate of increase into the nucleus following NE rupture for cells in D normalized by mobility. Error bars indicate ± SEM. Statistical significance: *, P<0.05; **<0.005; ***, P<0.0005 using an unpaired student t-test.

## Discussion

In these studies we have dissected multiple known functions and attributes of BAF as it responds to NE rupture and facilitates repair of the NE barrier. We found that BAF needs to be cytoplasmic, able to dimerize, and bind to DNA in a phosphoregulatable manner in order to prevent excessive leakage of contents between the nucleus and cytoplasm and to functionally reseal the NE permeability barrier. BAF does not rely on its binding to A-type lamins or, surprisingly, LEM domain proteins for it to prevent excessive leakage during a rupture or functionally reseal the NE barrier. Additionally, BAF appears capable of controlling the diffusion of cytosolic proteins into the nucleus during rupture based on their relative size, possibly by facilitating rapid membrane recruitment, physically impeding diffusion flow through the rupture hole via its targeting to the site of rupture, or both these processes.

In our initial studies we determined that, unlike untagged BAF, GFP-BAF is unable to rescue endogenous BAF’s role in response to NE rupture. This is not necessarily surprising given that GFP itself is larger than the BAF it is fused to, and it is commonly observed that the localization of GFP-BAF is substantially more NE-associated than endogenous BAF in mammalian cell lines (Shimi *et al*., 2004) suggesting that the behavior of GFP-BAF may not fully replicate that of the endogenous protein. However, we did find that cells lacking endogenous BAF and stably expressing GFP-BAF were viable without overt defects in postmitotic NE reformation or obvious changes in cell behavior, as has been previously reported in *C. elegans* (Gorjanacz *et al*., 2007b), suggesting that the GFP-BAF may at least partially rescue critical functions of BAF. It is also worth noting that we typically do not observe substantial defects in NE reformation or cell death in cells depleted of BAF by siRNA, even as long as 96 hr past initial knockdown. This suggests that, at least in the cell lines we examined, those phenotypes require longer time to occur and/or a more extensive BAF depletion, although the depletion is robust enough to cause substantial defects in its role in NE rupture. It was previously reported that GFP-BAF-WT but not GFP-BAF-L58R was able to rescue the loss of endogenous BAF during NE rupture in S-phase arrested U2OS cells depleted of lamin B1 (Young *et al*., 2020), whereas in our experiments GFP-BAF-WT is not functionally competent to rescue the loss of endogenous BAF but untagged L58R appears sufficient. We speculate that these differences might be attributed to experimental variables including the depletion of lamin B1, cell cycle arrest, NE rupture method, and/or cell-type.

We have developed and applied a new reporter, HSP90-GFP, to monitor nuclear rupture and repair that is independent of facilitated nuclear transport. We have yet to encounter a situation where the GFP-NLS does not function as a reliable reporter for the occurrence of NE rupture, barring a complete collapse of the Ran gradient with ATP depletion, but the extent of compartmental leakage and timing and rate of repair can be significantly impacted by physiologically relevant changes in nuclear transport (Andreu et al., 2022; Scott *et al*., manuscript in preparation). This is especially critical when utilizing the laser-induced NE rupture experiments that enable more control and consistency in the timing and extent of NE rupture events. Spontaneous ruptures, while more physiologically relevant, have unpredictable timing and in our experience only frequently occur in cancer cells and/or cell lines where the nuclear lamina is defective (De Vos *et al*., 2011; Yang *et al*., 2017; Chen *et al*., 2019; Janssen *et al*., 2022b), and therefore make a difficult model for quantifying the rupture leakage with high spatiotemporal resolution. Non-cancerous cell lines, such as those cells utilized in our studies, appear highly resistant to mechanical rupturing by constrained migration, and the BJ5ta fibroblasts are even resistant to confinement induced nuclear rupture (data not shown).

In BJ5ta cells, elevating levels of BAF has no appreciable impact on the rate of rupture leakage nor does it reduce the time to repair, suggesting that the endogenous level of BAF is sufficient to initiate repair and restore the nucleo-cytoplasmic barrier. However, progressively depleting endogenous BAF results in a progressive increase in rupture leakage and defective repair. This suggests that BAF doesn’t have a critical threshold to prevent leakage but instead functions in a dosage-dependent manner. However, functional repair of the NE is logically likely to be binary as the NE membranes are either repaired or not. It should not be a surprise that cytosol-confined BAF was capable of functionally rescuing the loss of endogenous BAF in repair of NE ruptures, given that this is the population of BAF that predominantly localizes to ruptures (Halfmann *et al*., 2019). Conversely, that nuclear-confined BAF was incapable of rescuing endogenous BAF may be unexpected since the majority of the endogenous BAF appears to be nuclear. BAF’s ability to not only bind DNA, but also to have that DNA binding be regulated by compartmental phosphorylation is key to its role in responding to NE ruptures. Non-phosphorylated BAF in the cytoplasm is primed to bind to dsDNA either from an exogenous source like DNA transfection or a virus, or from cytoplasmic exposure of genomic DNA following NE rupture. If this DNA binding is inhibited, either by mutation of a binding site residue (K6A) or phosphomimetic mutations of the N-terminus, the protein is unable to prevent rupture leakage or mediate NE repair. Yet if the DNA binding is unregulated by mutation of BAF’s N-terminal residues to prevent inhibitory phosphorylation, the protein becomes trapped in the nucleus, largely immobilized on the genomic DNA, and unable to participate in the response to rupture. If this protein can be partially shifted to the cytoplasm by addition of an NES, it regains functionality in preventing rupture leakage and enabling repair. Collectively, these results suggest that only a nominal amount of BAF capable of binding to DNA and located in the cytoplasm is necessary for this function.

BAF’s ability to interact and recruit transmembrane LEM-domain proteins was reported to be a critical component of NE rupture repair, as it was responsible for recruiting membrane to the rupture site, as well as the LEMD2-mediated recruitment of the ESCRT-III complex for membrane resealing (Halfmann *et al*., 2019; Young *et al*., 2020). Surprisingly, we have observed that a BAF variant inhibited in binding LEM-domain proteins (L58R) is capable of facilitating functional repair, raising the possibility that recruitment of LEM domain proteins to nuclear rupture sites may not be completely necessary for this process. It is possible that the previously reported repair defect from simultaneous loss of Ankle2, LEMD2, and Emerin (Halfmann *et al*., 2019) actually resulted from an alternate mechanism(s) caused by loss of these proteins from the cell as opposed to preventing their NE retention or recruitment. We also found that the BAF-A-type lamin interaction is dispensable for BAF’s role in reducing leakage and promoting repair of NE ruptures. BAF-A12T clearly prevents A-type lamin recruitment to ruptures, as has been previously reported (Janssen *et al*., 2022a; Sears and Roux, 2022), but failed to appreciably alter the reporter leakage or functional repair. It is possible that the absence of A-type lamins at these repaired ruptures make the nuclei prone to re-rupture due to a lack of stabilizing lamina at these sites, as has been reported (Janssen *et al*., 2022a), but this was not observed in our studies utilizing laser-induced ruptures.

Based on our BAF-A12T and BAF-L58R studies, BAF homodimerization is also a necessary attribute for control of rupture leakage and NE repair and appears to regulate these processes via a mechanism that is likely independent of lamin or LEM-domain recruitment. It is tempting to hypothesize that aside from facilitating membrane resealing, BAF enrichment at rupture sites may act to slow or block the diffusion of molecules in a size-dependent manner between the nuclear and cytosolic compartments, and may at least partially re-establish a nucleo-cytoplasmic barrier without complete resealing. In this scenario, it is possible that BAF could function at rupture sites to form a transient diffusion barrier for larger molecules, akin to a clot, through chromatin compaction (which BAF homodimerization would establish) and/or recruitment of additional proteins, as has been previously suggested (King and Lusk, 2019; Halfmann and Roux, 2021). Indeed, BAF was previously shown to bind and stiffen the surface of chromatin to form a size-dependent permeability barrier properties that were proposed to facilitate reformation of the NE after mitosis (Samwer *et al*., 2017). These same properties may function to reduce leakage and facilitate repair following NE rupture. The process of NE rupture repair appears to recapitulate NE reformation during open mitosis, however it remains unclear whether BAF’s evolution was driven by the need to reseal the NE following mitosis or to repair rupture of the interphase NE.

## Materials and Methods

### Cell Culture

BJ-5ta (ATCC CRL-4001), NIH3T3 (CRL-1658), and HEK293 Phoenix cell lines (National Gene Vector Biorepository) were cultured in Dulbecco’s modified Eagle medium with 4.5 g/L glucose, L-glutamine, and sodium pyruvate (DMEM; Corning). All media was supplemented with 10% (v/v) fetal bovine serum (FBS; Hyclone) at 37°C with 5% CO_2_ in a humidified incubator.

### Plasmids

All plasmids used in this study were generated using the Infusion cloning system (Takara) with primers containing 5’ flanking regions (15bp) complementary to the free-ends of the cloning vector and verified by sequencing. Enhanced GFP was used in all GFP-tagged constructs. GFP-BAF pBabe puro was constructed previously (Halfmann *et al*., 2019), and mCherry-BAF pBabe neo was constructed by PCR amplifying human BAF from GFP-BAF pBabe puro and recombining into BspEI-SalI cut mCherry-NLS pBabe neo (Halfmann *et al*., 2019). An siRNA-resistant human BAF was synthetically constructed (Gene Universal) by codon-optimizing human BAF (WT siRNA^R^ BAF) cDNA sequence using a codon-optimization tool (Integrated DNA Technologies; https://www.idtdna.com/CodonOpt) followed by verification of non-alignment with BAF siRNAs (Supplemental Table S1). This synthetic BAF transgene was then PCR amplified and cloned into IRES-GFP-NLS pBabe puro to create WT-BAF-IRES-GFP-NLS as previously described (Sears and Roux, 2022). All subsequent BAF variants were created using WT-BAF as a template for PCR then recombining into SnaBI-PmeI cut WT-BAF-IRES-GFP-NLS pBabe puro. NES-BAF was created by adding an N-terminal NES (GNELALKLAGLDI) to WT-BAF using a forward primer. NLS-BAF was was created by adding an N-terminal NLS (PKKKRKV) to WT-BAF using a forward primer. BAF-L58R was created by incorporating an L58R point mutation to forward and reverse primers and using overlap extension PCR to amplify the full BAF fragment. The BAF-A12T mutation was created by incorporating the A12T point mutation into a forward primer. Hsp90-GFP was first constructed by PCR amplifying HA-Hsp90 (a gift from William Sessa, Addgene plasmid #22487) and recombining into GFP pBabe puro digested with EcoRI-XhoI. To create each BAF variant in IRES-Hsp90-GFP pBabe puro, the GFP-NLS in each -IRES-GFP-NLS pBabe puro vector containing each BAF variant was removed by digesting with PmeI-SalI and replaced with Hsp90-GFP using in-fusion cloning. Empty-IRES-GFP-NLS and Empty-IRES-Hsp90-GFP pBabe puro was created by amplifing the MCS from pBabe puro and recombining into SnaBI-EcoRI digested WT-BAF-IRES-GFP-NLS and WT-BAF-IRES-Hsp90-GFP pBabe puro. Catalytically-inactive human cGAS (E225A/D227) –mCherry (a generous gift from Jan Lammerding) was cloned into a pBabe neo vector as previously described (Sears and Roux, 2022). 3GFP-Nup153 in pBabe puro was constructed by replacing N144 in 3GFP-N144 pCDNA3.1 (Birendra et al., 2017) with Nup153 using XhoI and AflII restriction enzyme sites. 3GFP-Nup153 was then PCR amplified and inserted into pBabe puro using EcoRI-SalI cut pBabe puro. GFP-Sec61β in pBabe puro (a generous gift from Indra Chandrasekar) was created by PCR amplifying Sec61β (a gift from Gia Voeltz, Addgene plasmid #49154) and recombining into XhoI-SalI cut GFP pBabe puro. The plasmid GFP-α-tubulin pBabe puro was created by PCR amplifying mouse α-tubulin from a cDNA template (a generous gift from Indra Chandrasekar) and recombining into XhoI-SalI cut GFP pBabe puro. All sequence data available upon request.

### Construction of stable cell lines

BJ5ta and NIH3T3 cell lines stably expressing fluorescent proteins were generated using retroviral transduction. For this, HEK293T Phoenix cells were seeded in 2 mL DMEM in a 6-well plate at 90% confluence and incubated overnight to allow for cell attachment. 1µg of pBabe-puro or -neo plasmid DNA encoding the protein of interest was transfected into the attached Phoenix cells using Lipofectamine 3000 (ThermoFisher) following manufacturer’s instructions. After an overnight incubation, cells were transferred to 32°C for 24 hr. The culture media was collected and filtered through a 0.45µm filter and added to BJ5ta cells (target cells), along with polybrene (2.5 µg/mL; Santa Cruz Biotechnology), and incubated at 37°C for 24 hr. Cells were trypsinized, collected through centrifuging at 250xg for 5 min, and incubated in fresh DMEM containing puromycin (0.5 µg/mL; Thermofisher) for selection of viral integration for 2-3 days. Expression of fluorescent proteins in cell lines was verified using immunofluorescence and immunoblot analysis.

### siRNA transfection

All siRNA transfections were performed using RNAiMAX (ThermoFisher) according to manufacturer’s instructions. Cells were trypsinized and seeded into 12-well plates at 70% confluency, and incubated overnight to allow cells to adhere to the bottom of the well. ON-TARGETplus SMARTpool siRNAs (Dharmacon) against human BAF (L-011536-02), mouse BAF (LU-062803-01) and an NT control (D-001810-01) were used for all transfections (Supplemental Table S1). 10 pmol siRNAs (0.75 µL of a 20 µM stock solution in RNAse free H_2_O) were dissolved in 75 µL of 1x OptiMEM. 4.5 µL of RNAiMAX was diluted in 75 µL of 1x OptiMEM. Both solutions were combined, mixed by pipetting and incubated for 5 min. The combined solution was then added to the attached cells and incubated for 72-96 hr. For either knockdown timeframe, cells were split and transferred to 35 mm glass-plated fluorodishes (World Precision Instruments) and allowed to attach for 48 hr prior to imaging. For BAF titrations, each diluted BAF siRNA was supplemented with NT Control siRNA to bring the final amount to 10pmol total siRNA, where applicable. Cell lysates were collected from fluorodishes after experiments and efficiency of knockdown was verified using immunoblotting or immunofluorescence with a BAF antibody.

### Immunofluorescence

Cells were grown on glass coverslips and fixed in 3% (wt/vol) paraformaldehyde/phosphate buffered saline (PFA) for 10 min and permeabilized by 0.4% (wt/vol) Triton X-100/PBS (PBST) for 15 min. Cells were labeled for 1 hr with rabbit anti-BAF (1:100; ab129184; Abcam), rabbit anti-Emerin (1:100; ab156871; Abcam) or rabbit anti-phospho-Lamin A/C (pSer22; 1:100; 2026S; Cell Signaling). Primary antibodies were detected using Alexa-Flour 488 conjugated goat anti-rabbit (1:1000, A21235, Thermofisher) and Hoescht dye 33342 to detect DNA. Coverslips were mounted using 10% (wt/vol) Mowiol 4–88 (Polysciences). Epifluorescent images were captured using a Nikon Eclipse NiE (40×/0.75 Plan Fluor Nikon objective; 40x/0.75 Plan Apo Nikon objective) microscope at room temperature with a CCD camera (CoolSnap HQ; Photometrics) linked to a workstation running NIS-Elements software (Nikon Melville, NY). All images were processed in Adobe Photoshop CS6 for cropping and brightness/contrast adjustment where applicable.

### Immunoblotting

Protein expression was analyzed from total cell lysate. 1.2 × 10^6^ cells were lysed in SDS/PAGE sample buffer (200 µL 2x sample buffer, 245 µL H_2_O, 50 µL SDS (20% w/v), and 5 µL DTT), boiled for 5 min, and sonicated to shear DNA. Proteins were separated on 4–20% gradient gels (Mini-PROTEAN TGX; Bio-Rad, Hercules, CA) and transferred to nitrocellulose membrane (Bio-Rad). Membranes were blocked with 10% (vol/vol) adult bovine serum and 0.2% Triton X-100 in PBS for 30 min, then incubated with appropriate primary antibodies. Primary rabbit anti-BAF (ab129184; Abcam) was used at a dilution of 1:1000. Rabbit anti-GFP (ab290; Abcam) was used at a dilution of 1:10000. Mouse monoclonal anti-tubulin was used as a loading control (sc-32293; Santa Cruz Biotechnology) at 1:2000 dilution. Primary antibodies were detected using HRP-conjugated anti-rabbit (1:5000; G21234; ThermoFisher), or anti-mouse (1:5000; F21453; ThermoFisher) antibodies. The signals from antibodies were detected using enhanced chemiluminescence via BioRad ChemiDoc™ MP Imaging System (BioRad, Hercules, CA).

### Laser-induced nuclear rupture and live cell imaging

Live cells expressing fluorescently-tagged proteins of interest were seeded on 35 mm glass-bottomed fluorodishes in pre-warmed phenol red-free DMEM with HEPES and FBS for imaging (Gibco). Cells were imaged on an Olympus FV1000 confocal microscope and FV10-ASW v4.1 software, with a temperature-controlled chamber set at 37°C and 60x/NA 1.42 PlanApo N oil immersion objective (3x zoom). The 488nm and 543nm scanning lasers were used for GFP and mCherry imaging. Laser-induced rupturing was carried out with a 405 nm excitation laser (8 sec for BJ5ta cells, 6 sec for NIH3T3 cells) at 100% power (tornado scan mode), while utilizing the SIM-scan feature for simultaneously imaging and laser rupturing. GFP photobleaching was performed using the main scanner and 488nm laser at 100% power for 30 sec. For testing the extent of nuclear rupture repair, cytoplasmic GFP-NLS or nuclear Hsp90-GFP was photobleached using the main scanner and 488nm laser at 100% power for 10 sec and re-imaged for an additional 4 minutes to observe any continued GFP-NLS or Hsp90-GFP leakage into the cytoplasm or nucleus, respectively. Laser-induced ruptures in cells expressing GFP-Nup153 and mCherry BAF were performed by focusing the 405nm laser on the apical surface of NE to observe rupture formation at NPCs. All GFP-NLS and Hsp90-GFP measurements were performed in ImageJ v1.52i by measuring the mean intensity of a region of interest (ROI) over time. The width of rupture holes using GFP-Sec61β was performed in ImageJ by manually setting an intensity threshold along the NE and measuring the distance of the rupture gap at the indicated time points. All images were processed in Adobe Photoshop CS6 for cropping and brightness/contrast adjustment when applicable.

### FLIP imaging

FLIP experiments were performed by continuously bleaching an area in the cytoplasm of Hsp90-GFP and GFP-α-tubulin expressing cells and measuring the loss of fluorescence from the cytoplasm distal from the bleach area. A non-bleached cell in the same frame was used as a control for photobleaching. The ratio of loss from the nucleus to the loss from the nucleus of the non-bleached cells was used to find the relative fluorescence loss. Significance was calculated by finding the rate of fluorescence lost over the first five minutes of photobleaching.

### Statistical analysis and Illustration design

Unpaired t-test, one way ANOVA with Dunnett’s post hoc multiple comparison test, or one way ANOVA with Tukey’s post hoc multiple comparison test were used to find significance, as denoted in the figure legends. Outliers were identified prior to analysis using ROUT analysis with Q set to 1%. All graphs represent mean values ± SEM (error bars). Statistical analysis, outlier identification, and graph generation were performed in GraphPad Prism v.7.02 (GraphPad Software Inc., San Diego, CA, USA). All illustrations were created using Biorender.com

## Acknowledgments

This work was supported by R35GM126949 to K.J.R. from the National Institutes of Health. The Imaging Core at Sanford Research, which facilitated these studies, is supported by Institutional Development Awards from the National Institute of General Medical Sciences and the National Institutes of Health under grant 1P30GM145398.

The authors declare no competing financial interests.

## Author contributions

Kyle J. Roux and Charles T. Halfmann contributed to conceptualization and experimental design. Kyle J. Roux contributed to project administration, supervision, and funding acquisition. Charles T. Halfmann, Kelsey L. Scott, and Rhiannon M. Sears performed experiments. Charles T. Halfmann, Kelsey L. Scott, and Rhiannon M. Sears performed formal data analysis. Kyle J. Roux and Charles T. Halfmann contributed to writing the original draft. Kyle J. Roux, Charles T. Halfmann, Kelsey L. Scott, and Rhiannon M. Sears performed draft review and editing.

