## Supplemental data for "Mechanisms by which barrier-to-autointegration factor regulates dynamics of nucleocytoplasmic leakage and membrane repair following nuclear envelope rupture"

**Table S1.** siRNAs used in this study

| Target | Species | Catalog No. | Name | Sequence (5' to 3') |
|---|---|---|---|---|
| BAF | Mouse | LU-062803-01 | J-062803-10 | UUGGUGACGUCCUGAGCAA |
| | | | J-062803-11 | CGAGAAUGGCUGAAGGAUA |
| | | | J-062803-12 | CCUAUGUACUCCAGGGUCU |
| BAF | Human | L-011536-02 | J-011536-10 | UACGACAAUAGCAAUCUUU |
| | | | J-011536-11 | UUCAUCUUUCUUUAGCACC |
| | | | J-011536-12 | UGCCACGAAGUCUCGGUGC |
| | | | J-011536-13 | UUGCCCAGGACUUCACCAA |
| NT Control | | D-001810-01 | J-017941-17 | CUAAAAUAUCGGUGGCGAA |
| | | | D-001810-02 | UGGUUUACAUGUUGUGUGA |
| | | | D-001810-03 | UGGUUUACAUGUUUUCUGA |
| | | | D-001810-04 | UGGUUUACAUGUUUUCCUA |

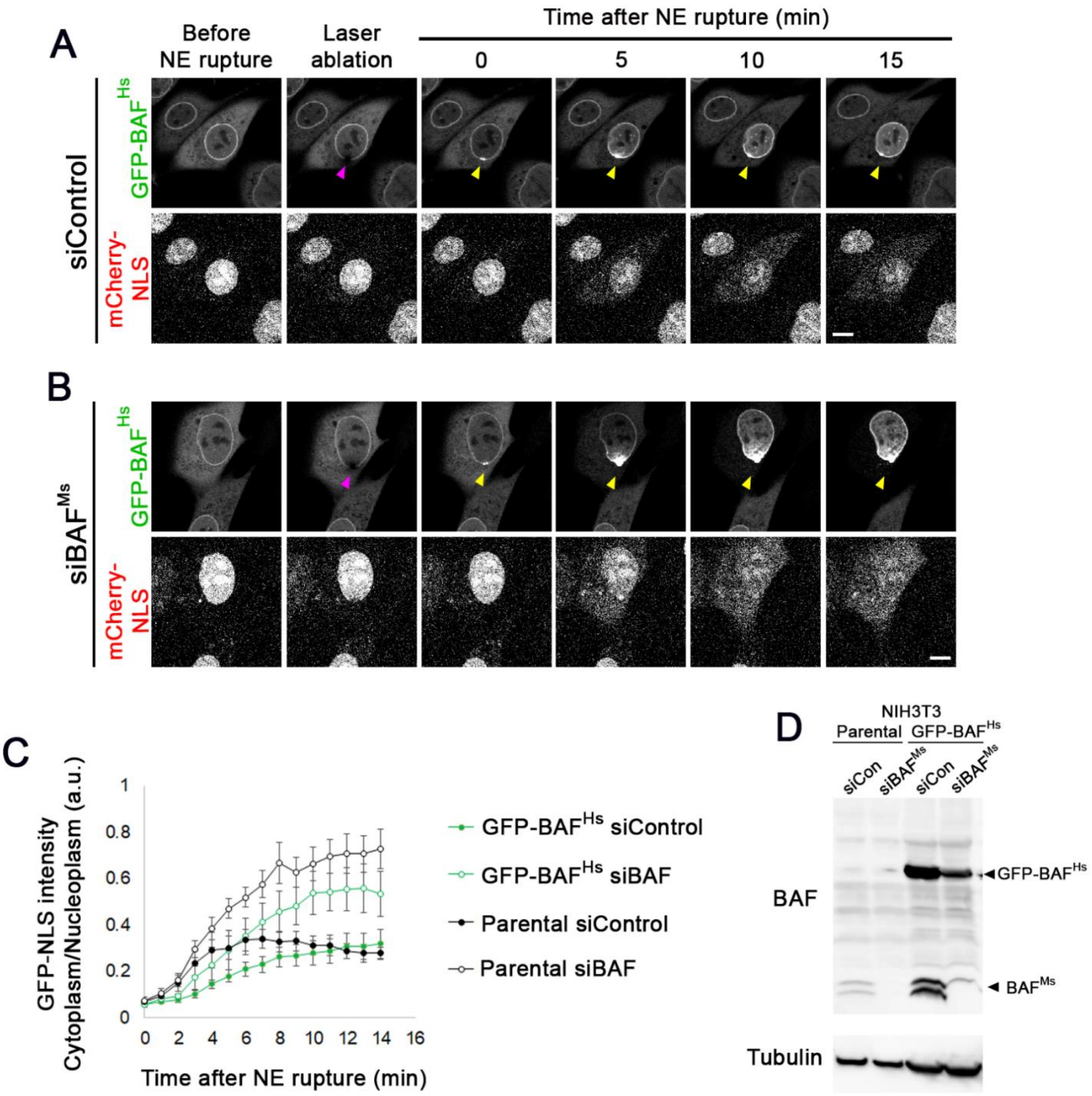


**Figure S1:** GFP-BAF does not functionally rescue the loss of endogenous BAF in the response to NE rupture. Sequential images of representative NIH3T3 cells expressing human GFP-BAF (GFP-BAF$^{Hs}$) and mCherry-NLS transfected with control siRNAs (A) or siRNAs against endogenous mouse BAF (siBAF$^{Ms}$, B) following laser-induced NE rupture. Purple arrow indicates site of laser ablation and yellow arrow indicates the site of GFP-BAF accumulation. Scale bar, 10µm. B) Cytoplasmic/nucleoplasmic ratios of mCherry-NLS measured over 15 min following laser-induced NE rupture from cells in A, from triplicate experiments (parental siControl, n=13; parental siBAF, n=12; GFP-BAF$^{Hs}$ siControl, n= 10; GFP-BAF$^{Hs}$ siBAF, n=8). C) Representative western blot of cell lysates from cells in A probed with BAF antibody, showing a loss of endogenous BAF but persistence of GFP-BAF when treated with siBAF$^{Ms}$. Scale bar, 10µm.

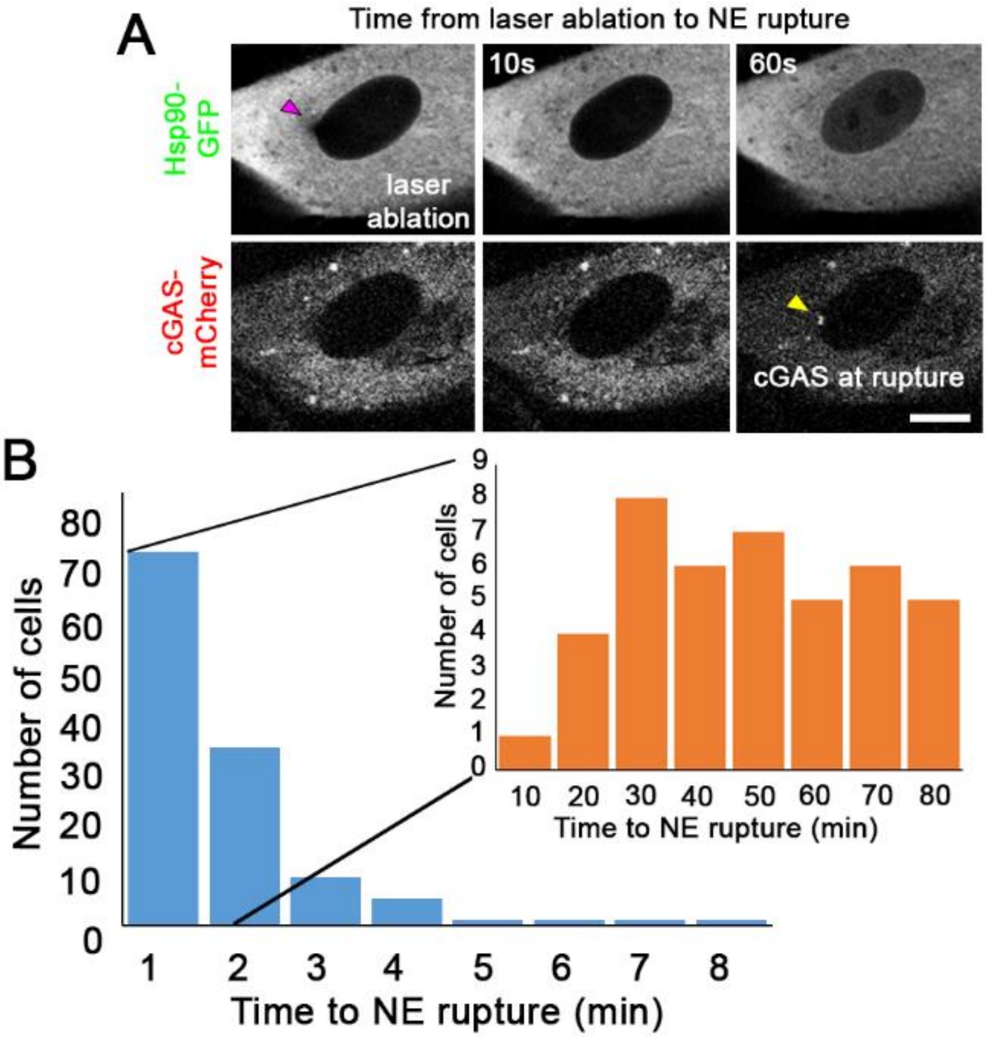


**Figure S2:** Timing of NE rupture following NE laser ablation. A) Sequential images of representative BJ5ta cells expressing Hsp90-GFP and cGAS mCherry following laser-ablation. Purple arrow indicates site of laser ablation on NE, yellow arrow shows rupture indicated by cGAS-mCherry enrichment. Scale bar: 10µm. B) Histogram of proportion of ruptures that occur after the laser ablation at the indicated time points, binned to every 1 minute. The insert histogram indicates proportion of ruptures that occur after the laser ablation during the first 80 sec, binned to every 10 sec *(n* values: 120 cells from 21 experiments).

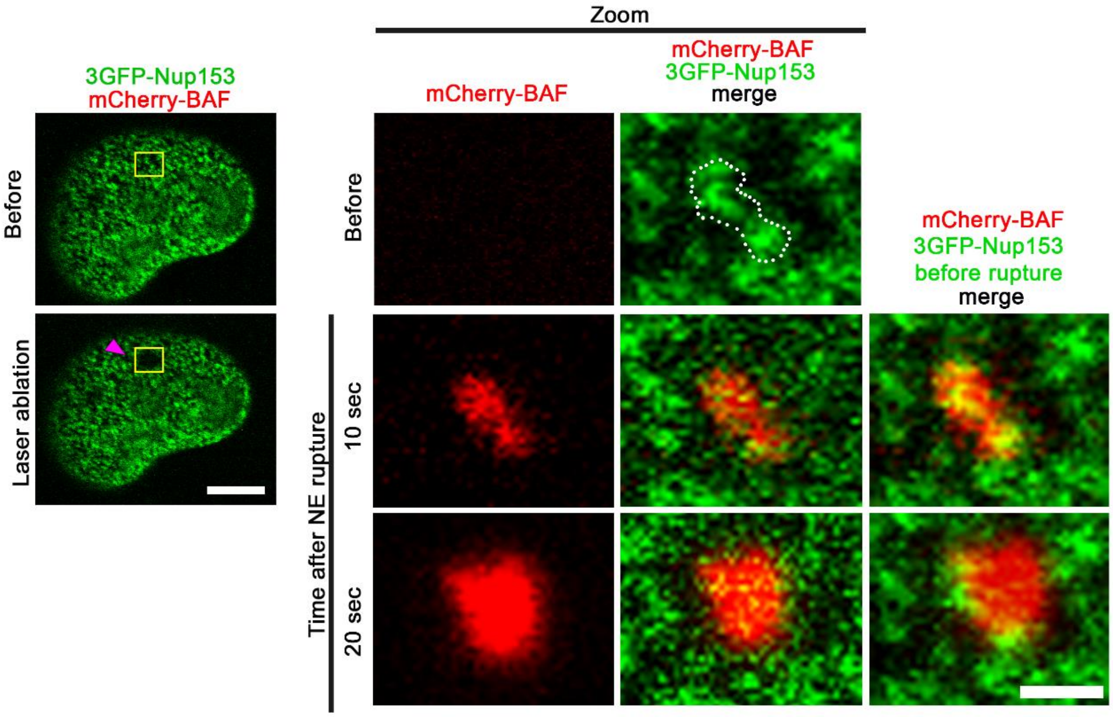


**Figure S3:** Laser-induced NE ruptures occur coincident with NPCs. Sequential images of a BJ5ta cell expressing a dynamic NPC marker (3GFP-Nup153) and mCherry-BAF during laser-induced NE rupture on the apical surface of the NE. Yellow box indicates zoomed in area on NE. Purple area indicates area of laser rupture. White outline indicates cluster of NPCs that enriches with mCherry BAF upon rupture, which then spreads and is contained within an NPC-free area on the envelope. Scale bar: 5 µm; zoomed image, 1µm.

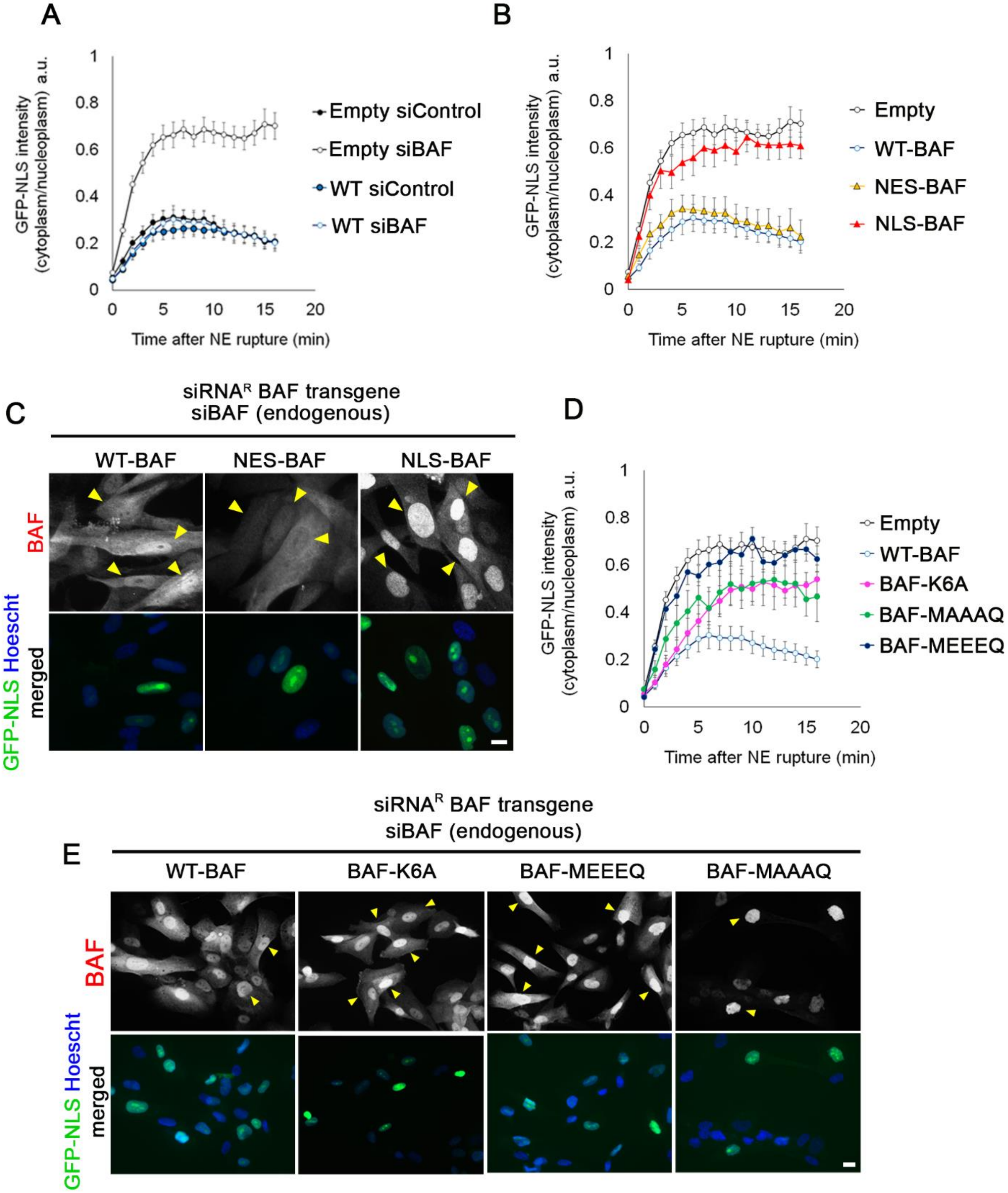

A
GFP-NLS intensity (cytoplasm/nucleoplasm) a.u.
Time after NE rupture (min)
Empty siControl
Empty siBAF
WT siControl
WT siBAF
B
GFP-NLS intensity (cytoplasm/nucleoplasm) a.u.
Time after NE rupture (min)
Empty
WT-BAF
NES-BAF
NLS-BAF
C
siRNA^R BAF transgene
siBAF (endogenous)
WT-BAF
NES-BAF
NLS-BAF
BAF
GFP-NLS Hoescht merged
D
GFP-NLS intensity (cytoplasm/nucleoplasm) a.u.
Time after NE rupture (min)
Empty
WT-BAF
BAF-K6A
BAF-MAAAQ
BAF-MEEEQ
E
siRNA^R BAF transgene
siBAF (endogenous)
WT-BAF
BAF-K6A
BAF-MEEEQ
BAF-MAAAQ
BAF
GFP-NLS Hoescht merged

**Figure S4:** Characterization and GFP-NLS ruptures of BAF variants affecting localization, DNA binding and/or phosphorylation. A) Cytoplasmic-to-nucleoplasmic ratios of GFP-NLS in BJ5ta cells expressing empty vector (empty) or WT-BAF (WT) IRES-GFP-NLS and treated with either siControl or siBAF for 96 hr prior to laser-induced NE rupture. Graph represents mean values ± SEM of the following number of cells: Empty siControl, n=21; Empty siBAF, n=13; WT siControl, n=16; WT siBAF, n=17 from triplicate experiments. B) Cytoplasmic-to-nucleoplasmic ratios of GFP-NLS after laser induced NE rupture in BJ5ta cells expressing empty-, WT-, NES-, or NLS-BAF and treated with siBAF for 96 hr prior to laser-induced NE rupture. Graph represents mean values ± SEM of the following number of cells: Empty, n=13; WT-BAF, n=17; NES-BAF, n=9; NLS-BAF, n=14 from at least duplicate experiments. C) Immunofluorescence images of BJ5ta cells expressing either WT-BAF, NES-BAF, or NLS-BAF and IRES-GFP-NLS treated with siBAF for 96 hr and probed with BAF antibody. Yellow arrows indicate expressing cells based on GFP-NLS expression. Scale bar; 10 µm. D) Cytoplasmic-to-nucleoplasmic ratios of GFP-NLS after laser induced NE rupture in BJ5ta cells expressing empty and BAF-WT, -K6A, -MEEEQ, and -MAAAQ and treated with siBAF for 96 hr prior to laser-induced NE rupture. Graph represents mean values ± SEM of the following number of cells: Empty, n=13; WT-BAF, n=17; BAF-K6A, n=10; BAF-MEEEQ, n=4, and BAF-MAAAQ n=5 from at least duplicate experiments. E) Immunofluorescence images of BJ5ta cells expressing either WT-BAF, BAF-K6A, BAF-MEEEQ or MAAAQ BAF with IRES-GFP-NLS treated with siBAF for 96 hr and probed with BAF antibody. Yellow arrows indicate expressing cells based on GFP-NLS expression. Scale bar; 10µm.

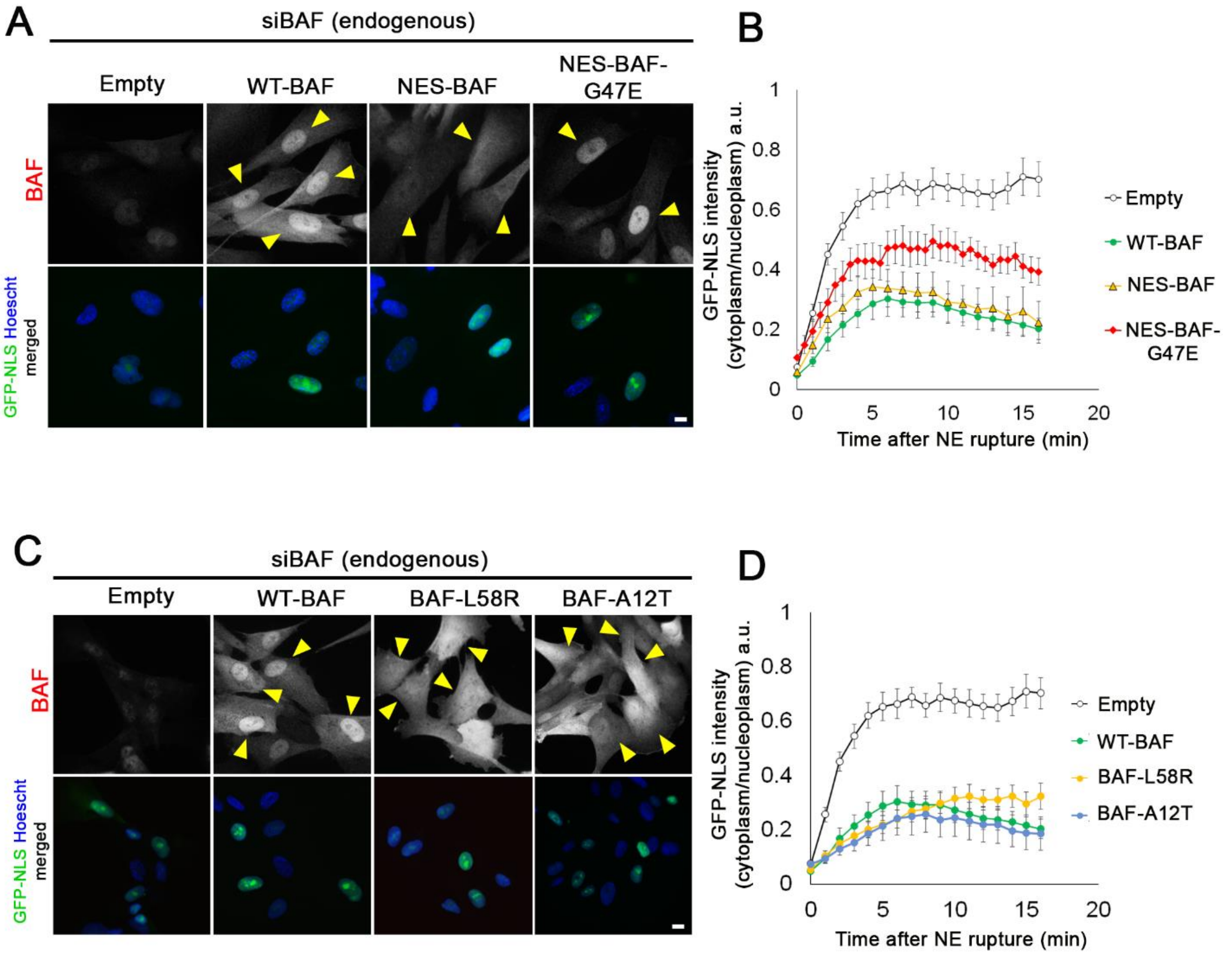


**Figure S5:** Characterization and GFP-NLS ruptures of BAF variants that alter homodimerization, LEM-domain protein binding, and lamin binding. A) Immunofluorescence images of BJ5ta cells expressing Empty, WT-BAF, NES-BAF, or NES-BAF-G47E and IRES-GFP-NLS treated with siBAF for 96 hr and probed with BAF antibody. Yellow arrows indicate expressing cells based on GFP-NLS expression. Scale bar; 10 µm. B) Cytoplasmic-to-nucleoplasmic ratios of GFP-NLS in BJ5ta cells expressing Empty, WT-BAF, NES-BAF, or NES-BAF-G47E and IRES-GFP-NLS and treated with siBAF for 96 hr prior to laser-induced NE rupture. Graph represents mean values ± SEM of the following number of cells: Empty, n=13; WT-BAF n=17, NES-BAF n= 9, NES-BAF-G47E n= 13 from at least duplicate experiments. C) Immunofluorescence images of BJ5ta cells expressing either Empty, WT-BAF, BAF-L58R, or BAF-A12T with IRES-GFP-NLS treated with siBAF for 96 hr and probed with BAF antibody. Yellow arrows indicate expressing cells based on GFP-NLS expression. Scale bar; 10µm. D) Cytoplasmic-to-nucleoplasmic ratios of GFP-NLS in BJ5ta cells expressing Empty, WT-BAF, BAF-L58R, or BAF-A12T and IRES-GFP-NLS and treated with siBAF for 96 hr prior to laser-induced NE rupture. Graph represents mean values ± SEM of the following number of cells: Empty, n=13; WT-BAF, n=17; BAF-L58R, n=13; BAF-A12T, n=10 from at least duplicate experiments.

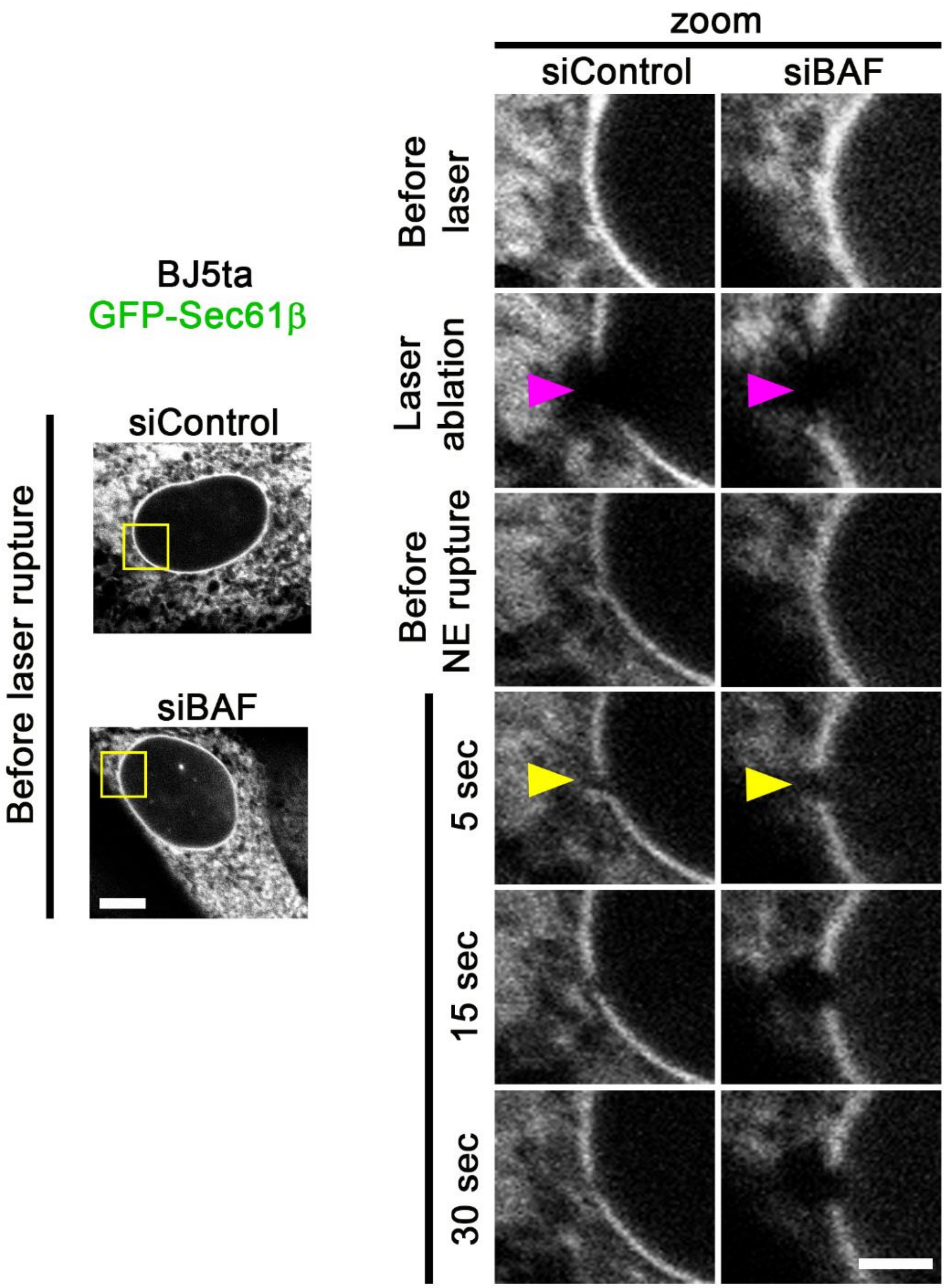


**Figure S6:** Rupture holes rapidly close in the presence of BAF. A) Representative images of BJ5ta cells expressing GFP-Sec61β treated with either siControl or siBAF and imaged to observe the width of the NE gap during the first 30 seconds following laser-induced NE rupture. Yellow box indicates zoomed-in area on the NE. Pink arrow indicates the site of laser ablation on the NE, yellow arrow indicates the site of rupture. Experiments were performed in triplicate with the following number of cells: siControl, n=15; siBAF, n=14. Scale bar; 10μm on full scale images (left), 1μm on zoomed images.
